# Short- and long-range roles of UNC-6/Netrin in dorsal-ventral axon guidance *in vivo* in *Caenorhabditis elegans*

**DOI:** 10.1101/2024.04.23.590737

**Authors:** Kelsey M. Hooper, Erik A. Lundquist

**Affiliations:** University of Kansas, Department of Molecular Biosciences, Program in Molecular, Cellular, and Developmental Biology

## Abstract

Recent studies in vertebrates and *Caenorhabditis elegans* have reshaped models of how the axon guidance cue UNC-6/Netrin functions in dorsal-ventral axon guidance, which was traditionally thought to form a ventral-to-dorsal concentration gradient that was actively sensed by growing axons. In the vertebrate spinal cord, floorplate Netrin1 was shown to be largely dispensable for ventral commissural growth. Rather, short range interactions with Netrin1 on the ventricular zone radial glial stem cells was shown to guide ventral commissural axon growth. In *C. elegans*, analysis of dorsally-migrating growth cones during outgrowth has shown that growth cone polarity of filopodial extension is separable from the extent of growth cone protrusion. Growth cones are first polarized by UNC-6/Netrin, and subsequent regulation of protrusion by UNC-6/Netrin is based on this earlier-established polarity (the Polarity/Protrusion model). In both cases, short-range or even haptotactic mechanisms are invoked: in vertebrate spinal cord, interactions of growth cones with radial glia expressing Netrin-1; and in *C. elegans*, a potential close-range interaction that polarizes the growth cone. To explore potential short-range and long-range functions of UNC-6/Netrin, a potentially membrane-anchored transmembrane UNC-6 (UNC-6(TM)) was generated by genome editing. *unc-6(tm)* was hypomorphic for dorsal VD/DD axon pathfinding, indicating that it retained some *unc-6* function. Polarity of VD growth cone filopodial protrusion was initially established in *unc-6(tm)*, but was lost as the growth cones migrated away from the *unc-6(tm)* source in the ventral nerve cord. In contrast, ventral guidance of the AVM and PVM axons was equally severe in *unc-6(tm)* and *unc-6(null)*. Together, these results suggest that *unc-6(tm)* retains short-range functions but lacks long-range functions. Finally, ectopic *unc-6(+)* expression from non-ventral sources could rescue dorsal and ventral guidance defects in *unc-6(tm)* and *unc-6(null)*. Thus, a ventral directional source of UNC-6 was not required for dorsal-ventral axon guidance, and UNC-6 can act as a permissive, not instructive, cue for dorsal-ventral axon guidance. Possibly, UNC-6 is a permissive signal that activates cell-intrinsic polarity; or UNC-6 acts with another signal that is required in a directional manner. In either case, the role of UNC-6 is to polarize the pro-protrusive activity of UNC-40/DCC in the direction of outgrowth.

## Introduction

UNC-6/Netrin is a conserved regulator of dorsal-ventral axon guidance (Hedgecock *et al.* 1990; Ishii *et al.* 1992; Serafini *et al.* 1994; Norris and Lundquist 2011)(reviewed in (Boyer and Gupton 2018)). It was previously thought that the UNC-40/DCC receptor mediated ventral growth and attraction to the Netrin source (Chan *et al.* 1996; Keino-Masu 1996; Kolodziej *et al.* 1996; Deiner *et al.* 1997), whereas the UNC-5 receptor mediated dorsal growth and repulsion from the Netrin source (Leung-Hagesteijn 1992; Hamelin *et al.* 1993). Recent studies of UNC-6/Netrin on the growth cones of axons *in vivo* in *C. elegans* show that UNC-40/DCC and UNC-5 are each involved in both ventral and dorsal guidance. In the HSN neuron, which extends an axon ventrally, the protrusive activity of UNC-40 is polarized ventrally by UNC-6, and refined and maintained by UNC-5 (the Statistically-Oriented Asymmetric Localization (SOAL) model) (Kulkarni *et al.* 2013; Yang *et al.* 2014; Limerick *et al.* 2017). In VD growth cones, which grow dorsally, UNC-6/netrin first polarizes the growth cone via UNC-5, such that UNC-40 protrusive activity is localized dorsally, and then maintains this polarity and regulates growth cone protrusion (Norris and Lundquist 2011; Norris *et al.* 2014; Gujar *et al.* 2018). UNC-6 drives protrusion dorsally through UNC-40, and inhibits protrusion ventrally and laterally through UNC-5, resulting in net dorsal outgrowth (the Polarity/Protrusion model) (Norris and Lundquist 2011; Norris *et al.* 2014; Gujar *et al.* 2018; Mahadik and Lundquist 2023b). Common to the SOAL and Polarity/Protrusion models is the polarization of pro-protrusive UNC-40/DCC activity by UNC-6 and UNC-5, which results in directed protrusion and growth cone advance. In ventral growth (SOAL), UNC-6 polarizes UNC-40 toward the ventral UNC-6/netrin source, and in dorsal growth, UNC-6 polarizes UNC-40 activity away from the ventral source.

At the time of VD growth cone outgrowth, UNC-6 is expressed in the VA and VB neurons in the ventral nerve cord, whose processes extend the length of the ventral nerve cord (Wadsworth *et al.* 1996). The dorsally-directed VD/DD motor axons also extend in the ventral nerve cord in proximity to the VA/VB processes. In contrast, the HSN and AVM/PVM neuron cell bodies reside laterally and extend processes ventrally toward the ventral nerve cord. Thus, UNC-6 might have both short- and long-range functions, the later relying on diffusion of UNC-6 away from its ventral expression source. In the vertebrate spinal cord, a long-range function of Netrin1 expression in the floorplate was invoked to explain ventral guidance of dorsal commissural axons (Kennedy *et al.* 1994; Serafini *et al.* 1996; Tessier-Lavigne and Goodman 1996). However, in *Drosophila*, Netrin-1 was shown to have close-range roles independent of DCC (Keleman and Dickson 2001). Further, cell-specific knock-out experiments showed that floorplate Netrin1 was largely dispensable for commissural axon guidance (Dominici *et al.* 2017; Varadarajan and Butler 2017; Yamauchi *et al.* 2017; Morales 2018). Instead, Netrin1 expression on the radial glial ependymal cells was required, in a short-range, possibly contact-mediated process. Thus, vertebrate Netrin1 might also have short-range functions.

In the Polarity/Protrusion hypothesis of VD dorsal growth cone guidance, a short-range interaction with UNC-6 in the ventral nerve cord polarizes the growth cone, and a longer-range role dependent upon diffusible UNC-6 maintains this polarity and regulates extent of protrusion biased dorsally (Norris and Lundquist 2011; Norris *et al.* 2014; Gujar *et al.* 2018; Mahadik and Lundquist 2023b). In this model, it is predicted that only the initial, short-range interaction is required to come from a directional source (ventral), whereas the long-range function is predicted to not require a directional source to maintain polarity and regulate protrusion.

To analyze potential short- and long-range roles of UNC-6/Netrin, the endogenous *unc-6* gene was edited such that a transmembrane domain was introduced at the C-terminus of the UNC-6 protein (the *unc-6(lq154)* genome edit, or *unc-6(tm)*). Transmembrane UNC-6 (UNC-6(TM)) is predicted to be anchored in the plasma membrane of the cell in which it is expressed. *unc-6(tm)* was hypomorphic, and animals were less severely uncoordinated and had fewer VD/DD axon guidance defects compared to the *unc-6(ev400)* null. VD growth cones were initially properly polarized in *unc-6(tm)*, but polarity was lost as the growth cones advanced dorsally away from the ventral nerve cord. In contrast *unc-6(null)* growth cones were unpolarized as soon as they emerged from the ventral nerve cord. This suggests that UNC-6(TM) is sufficient to initially polarize the VD growth cone (at short range) but that diffusible UNC-6 is required to maintain polarity (long-range). The ventral guidance of the AVM and PVM axons in *unc-6(tm)* resembled the *unc-6(null)*, consistent with *unc-6(tm)* lacking long-range UNC-6 activity.

The Polarity/Protrusion model predicts that maintenance of VD growth cone polarity would not require a directional source of UNC-6. To test this idea, *unc-6(+)* was expressed ectopically in the pharynx in the anterior and in dorsal body wall muscle. In a wild-type background, ectopic *unc-6(+)* expression had no effect on VD/DD axon guidance. Surprisingly, ectopic *unc-6(+)* expression strongly rescued VD/DD dorsal axon guidance defects and VD growth cone polarity defects of both *unc-6(tm)* and *unc-6(null)* mutants. Ectopic *unc-6(+)* expression also strongly rescued AVM/PVM and HSN ventral guidance defects in both mutants. These data indicate that a ventral directional source of UNC-6 is not required for dorsal-ventral axon guidance and growth cone polarity, and indicate that UNC-6 might act as a permissive rather than instructive signal in axon guidance.

### *unc-6(lq154)* can encode a transmembrane UNC-6 molecule

Cas9 genome editing was used to create *unc-6(lq154)* (also called *unc-6(tm)*), which has the potential to encode a transmembrane UNC-6 molecule (UNC-6(TM)) (Figure 1A). Intron 12 and exon 13 were replaced by a synthetic mini-intron and a recoded exon 13 that retained the identical coding potential of wild-type exon 13 but lacked the stop codon. *Green fluorescent protein (gfp)* coding region without a stop codon was fused in frame to exon 13. This was followed by a synthetic intron containing the *Hygromycin*-*resistance* (*HygR*) selection cassette in the opposite antiparallel direction, and a new exon containing the coding region for the transmembrane domain of the *cdh-3* gene fused in frame to *gfp*. The genomic sequence of *unc-6(lq154)* was confirmed by sequencing overlapping PCR products across the region. The sequence of the repair plasmid used to create *unc-6(lq154)*, which is the sequence of the *unc-6(lq154)* mutation, can be found in Supporting Information.

**Figure 1.**
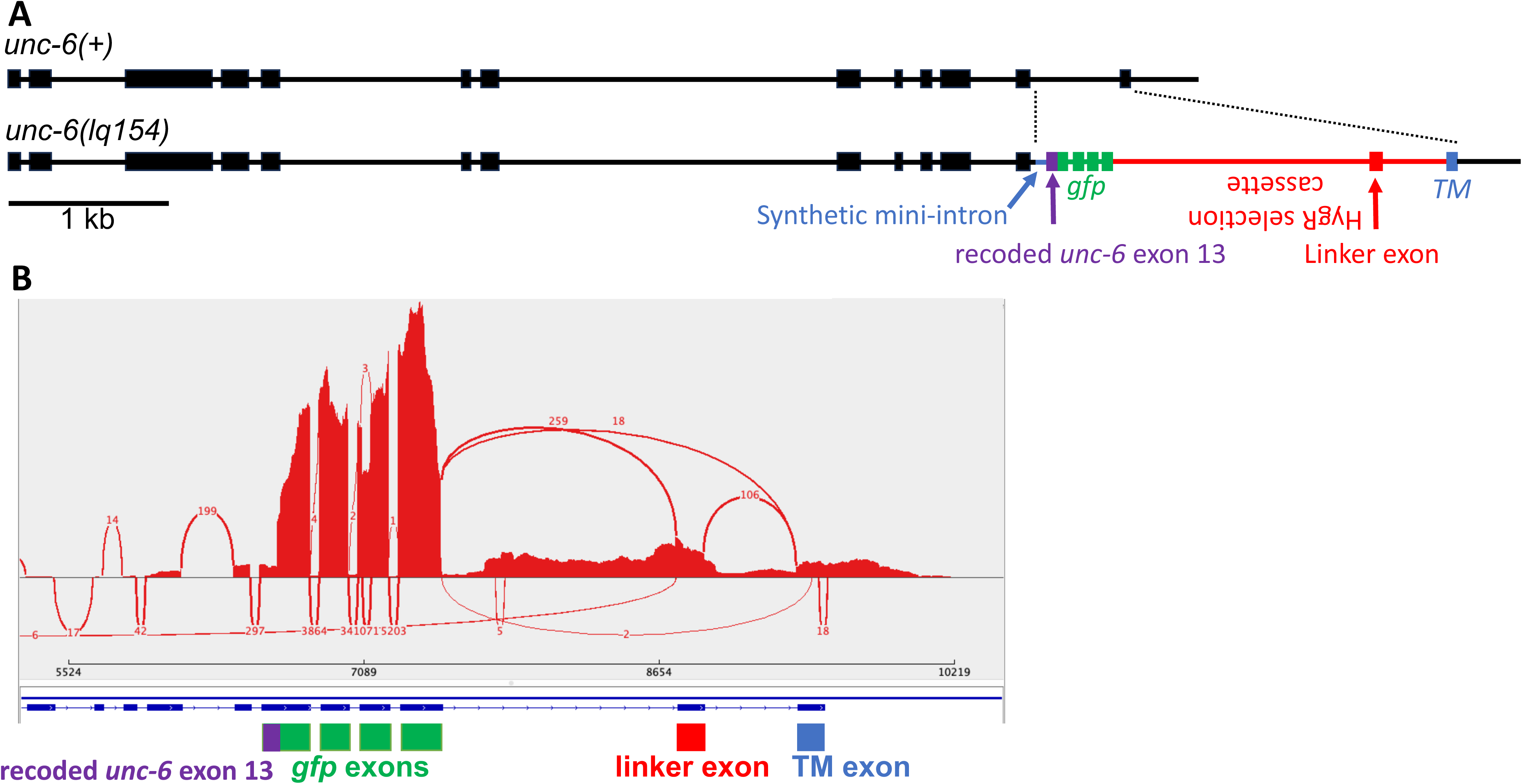
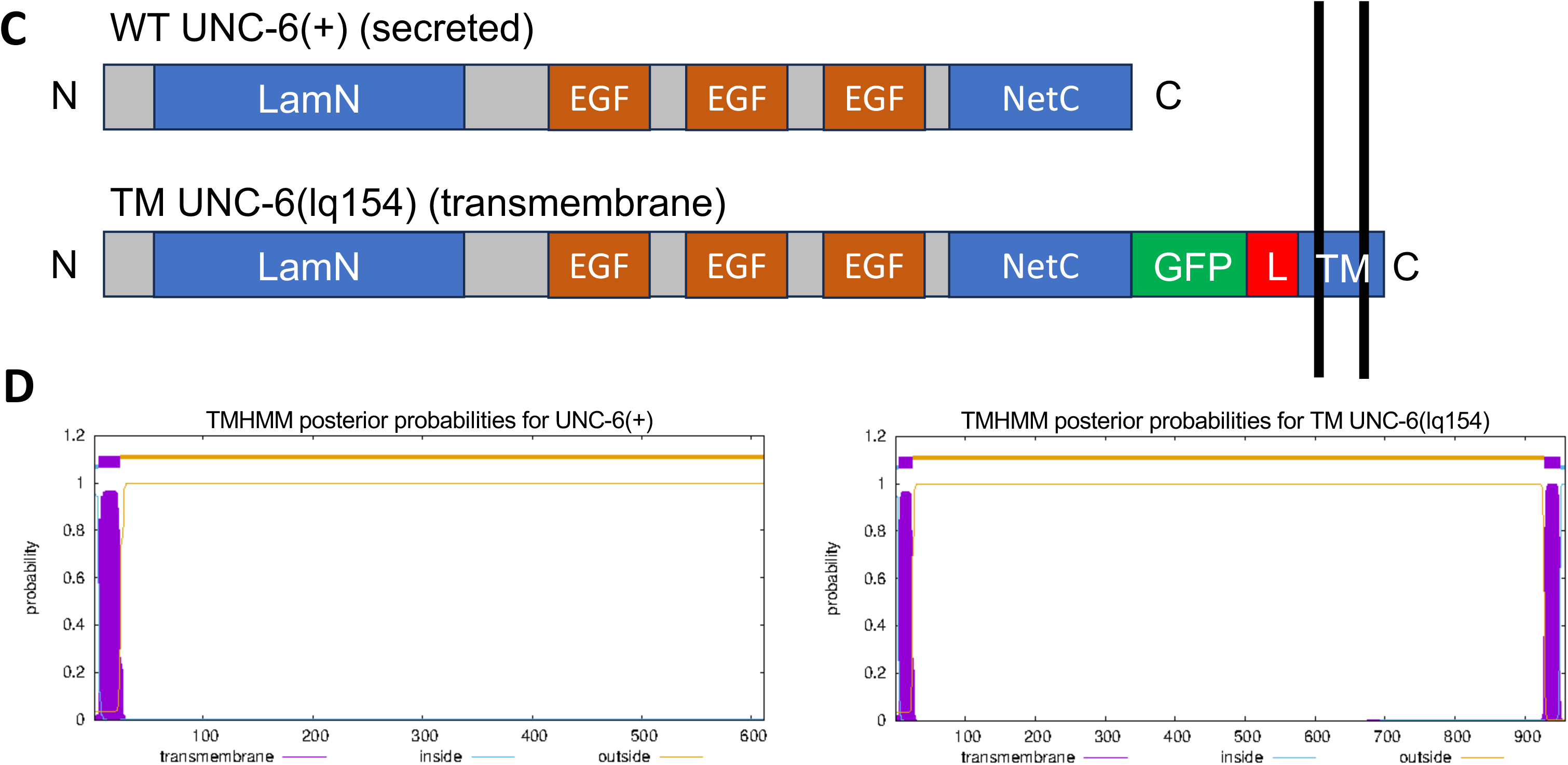
*unc-6(lq154)* genome edit structure. A) The *unc-6(+)* and *unc-6(lq154)* gene structures. Boxes represent exons and lines represent introns. In *unc-6(lq154)*, intron 12 and exon 13 (dashed line) were replaced by: a synthetic mini intron (blue), a recoded exon 13 lacking a stop codon but with the identical coding potential as *unc-6(+)* (purple), *green fluorescent protein* in frame with *unc-6* (green), a *Hygromycin-resistance* cassette, with the *HygR* gene on the opposite strand (red), an exon encoding the transmembrane domain (TM) from the *cdh-3* gene with a stop codon (blue), followed by the endogenous *unc-6* 3’ UTR region. B) A Sashimi plot from the Integrated Genome Viewer of RNA-seq reads from *unc-6(lq154)* aligned to a reference genome edited to include the *unc-6(lq154)* genome edit. The gfp to linker exon splice occurred 18 times, but the predominant splice variant (259) included a novel, likely artifactual, exon in the *HygR* cassette (the linker exon). This exon is 150-bp long and when translated, is in frame with *gfp* and the TM exon and contains no in-frame stop codons. The exon adds an additional 50 amino acid residues to the molecule with no significant similarity to other proteins. In many cases, the introns flanking the linker exon were not removed, resulting in in-frame premature stop codons in each case. Read coverage of *gfp* exons was greater than that of surrounding *unc-6(lq154)* sequence because the strain used for RNA-seq included the *Punc-25::*gfp transgene (*juIs76)*. C) A diagram of the UNC-6(+) and UNC-6(lq154) TM molecules. LamN is the laminin N domain; EGF are epidermal growth factor repeat domains; NetC is the Netrin C-terminal domain; GFP is green fluorescent protein; L is the linker exon; and TM is the transmembrane domain. UNC-6 is secreted, and UNC-6(lq154) is predicted to be a transmembrane protein. D) Trans Membrane Hidden Markov Model (TMHMM) analysis of UNC-6(+) predicts a secreted protein with an N-terminal signal peptide, and that UNC-6(lq154) encodes a transmembrane protein with an N-terminal signal peptide and a C-terminal transmembrane region.

The *HygR* gene was flanked by *LoxP* sites, but experiments to remove the *LoxP* region using *Cre* expressed in the germline were unsuccessful. To assess transcripts from the genome-edited region, RNA-seq from *unc-6(lq154)* was conducted. Reads were aligned to a reference genome file that was edited to include the *unc-6(lq154)* sequence. Reads from transcripts spanning the removal of the synthetic mini-intron and the *HygR* selection cassette intron were identified (Figure 1B). However, a common splicing pattern included an unpredicted novel exon (the linker exon) within the *HygR* cassette (Figure 1A and B). This linker exon is 150 bp in length and retains the reading frame from *gfp* to the transmembrane domain-containing final exon. Furthermore, it includes no in-frame stop codons. The net result of inclusion of the exon is an additional 50 amino acids between GFP and the transmembrane region with no detectable similarity to other proteins (Figure 1C). In many cases, the introns flanking the linker exon were not removed, resulting in in-frame premature stop codons in each case, and likely nonsense-mediated decay. This is evidenced by the *unc-6(lq154)* Unc and axon guidance phenotypes, which would not be expected if full-length UNC-6 without the transmembrane domain was being produced in *unc-6(lq154)*. In sum, *unc-6(lq154)* is predicted to encode transmembrane UNC-6 molecules with and without the 50 extra linker amino acids. The potential UNC-6(TM) protein is predicted to be a transmembrane protein using the Trans Membrane Hidden Markov Model (TMHMM) algorithm (Figure 1D).

### *unc-6(lq154tm)* is hypomorphic for dorsal VD/DD axon guidance

The *unc-6(ev400)* null mutants are severely uncoordinated and display severe defects in the dorsal to ventral guidance of the VD and DD GABA-ergic motor axons. The VD/DD cell bodies reside in the ventral nerve cord. Axons extend anteriorly in the ventral nerve cord, and then turn dorsally for commissural migration (Figure 2A and D). In wild-type, an average of 16 VD/DD ventral-to-dorsal commissures are apparent. In *unc-6(ev400)* null mutants, an average of 10 commissures showed any obvious extension at all from the ventral nerve cord (Figure 2F and G), an average of 3.2 crossed the lateral midline, (Figure 2H), and an average of fewer than one commissure per animal reached the dorsal nerve cord (Figure 2I).

**Figure 2.**
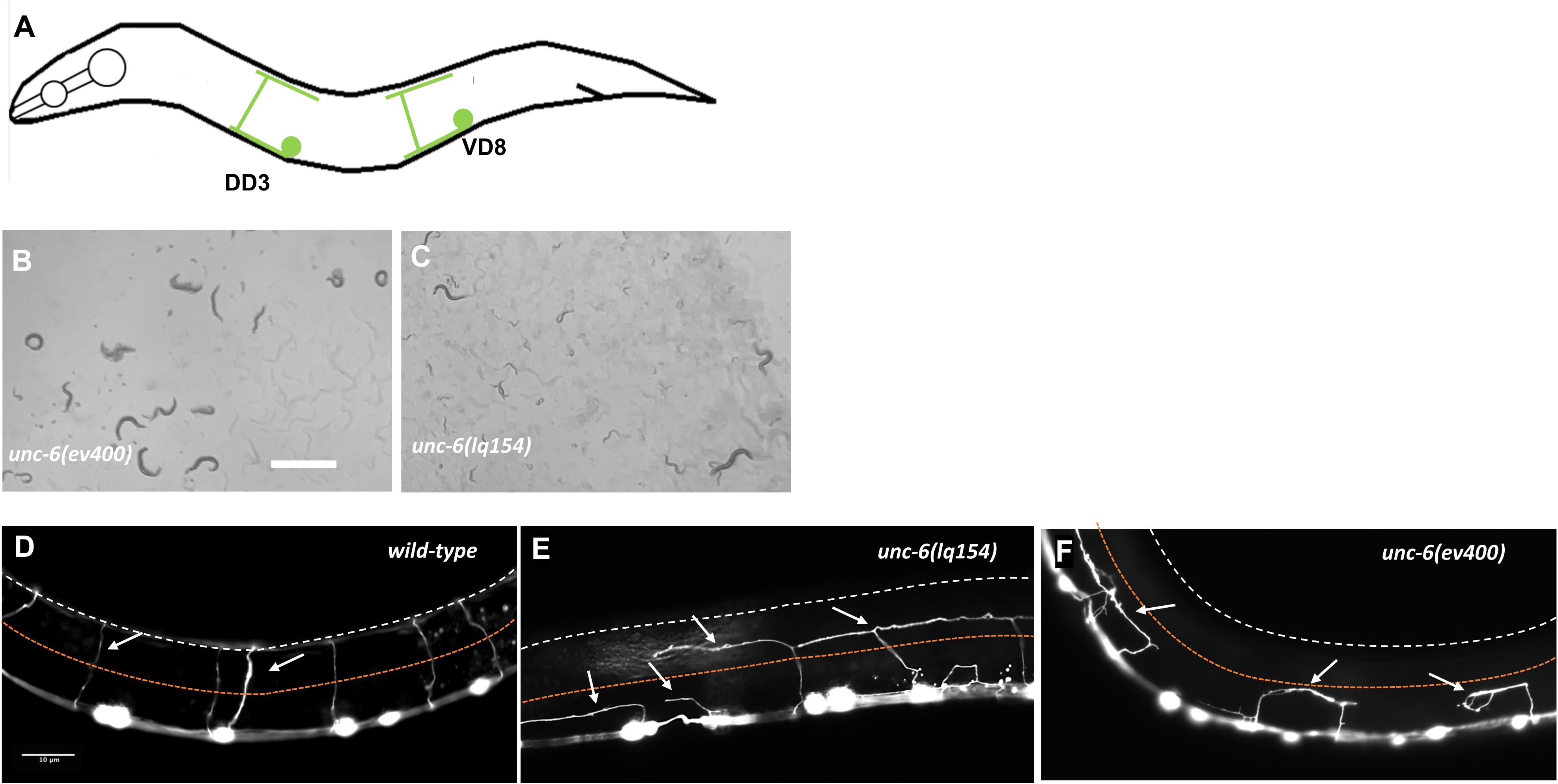
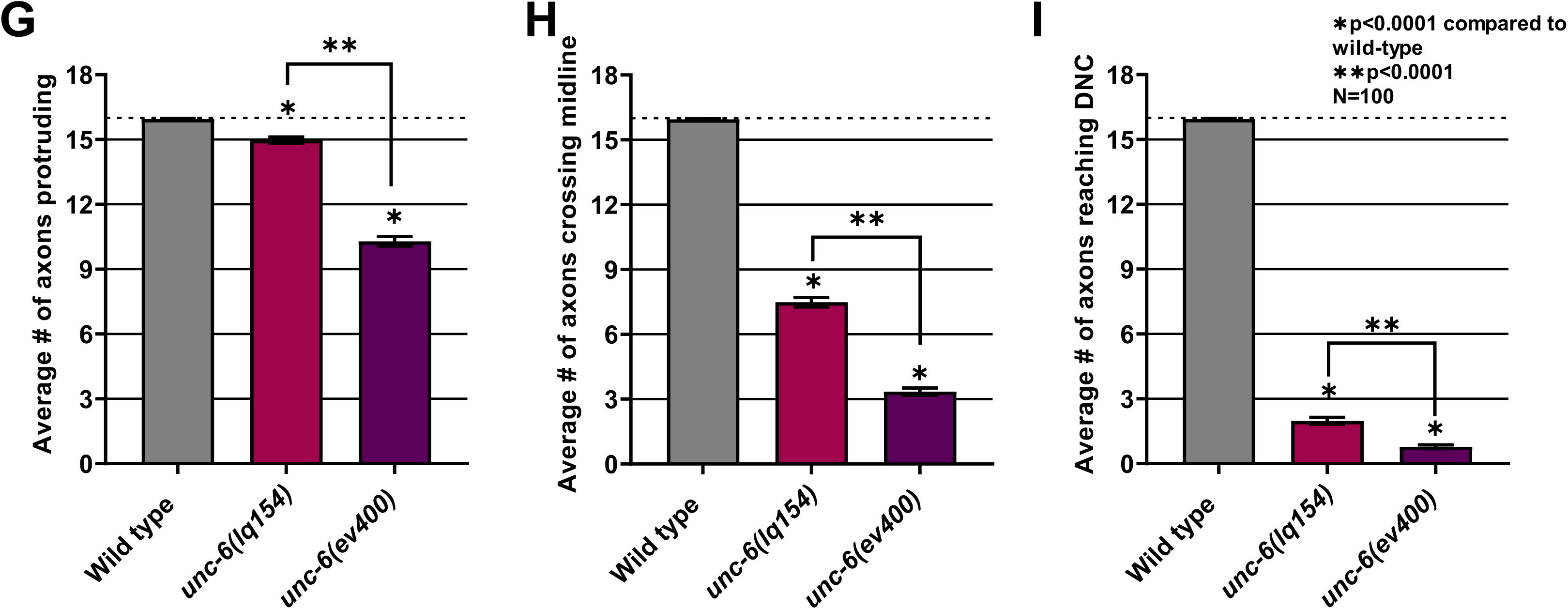
VD/DD axon guidance defects in *unc-6* mutants. A) A diagram showing the structure of the VD and DD axons. Anterior is to the left and dorsal is up. Only one of 13 VDs and one of 6 DDs is shown. Cell bodies reside in the ventral nerve cord. Axons extend anteriorly in the VNC, and then turn dorsally to migrate commissurally to the dorsal nerve cord, where they turn posteriorly and extend in the DNC, tiling the VNC and DNC. B,C) Light micrographs of *unc-6(ev400)* and *unc-6(lq154)* animals in lawns of *E. coli* on NGM plates. *unc-6(ev400)* animals are uncoordinated, with kinked body posture, and tend to not move and stay clustered in one place. *unc-6(lq154)* are less uncoordinated, show a more defined S-shaped body posture and movement, and tend the move and spread more evenly across the lawn of *E. coli*. The scale bar in B represents 1 mm. D-F) Fluorescent micrographs of VD/DD commissural axons in wild-type and *unc-6* mutants expressing the *juIs76[Punc-25::gfp]* transgene. The lateral midline is indicated by a dashed orange line and the dorsal nerve cord is indicated by a dashed white line. White arrows point to axons. The scale bar in D represents 10 μm for all images. G-I) Graphs quantifying VD/DD axon pathfinding defects in *unc-6* mutants. The X-axis represents genotype and the Y axis the number if commissures observed in animals. In wild-type, an average of 16 commissures extend from the ventral nerve cord to the dorsal nerve cord. Error bars represent standard error of the mean. Statistical significance was determined using a two-tailed *t*-test with unequal variance. G) The average number of commissures emanating from the ventral nerve cord. H) The average number of commissures extending past the lateral midline. I) The average number of commissures reaching the DNC.

*unc-6(lq154)* animals were less severely uncoordinated than *unc-6(ev400)* (Figure 2B and C). *unc-6(lq154)* displayed VD/DD axon guidance defects (Figure 2E), but significantly fewer compared to *unc-6(ev400)* null mutants: significantly more commissures emanated from the ventral nerve cord (15 compared to 10) (Figure 2G); significantly more crossed the lateral midline (7.8 compared to 3.2) (Figure 2H), and significantly more reached the dorsal nerve cord (2 compared to <1) (Figure 2I). These data indicate that *unc-6(lq154)* is not a complete loss of *unc-6* function and is hypomorphic for dorsal VD/DD axon guidance.

### *unc-6(lq154tm)* resembles the *unc-6* null phenotype for AVM/PVM ventral axon guidance

The AVM and PVM mechanosensory neurons reside laterally and are positioned dorsally away from the ventral nerve cord (Figure 3A, B, and E). AVM and PVM extend axons ventrally toward the ventral nerve cord. *unc-6(ev400)* null mutants display 72% and 73% failure of AVM and PVM ventral guidance, respectively (Figure 3C-I). The misguided axons extended anteriorly and failed to reach the ventral nerve cord. *unc-6(lq154)* AVM and PVM ventral guidance did not differ significantly from *unc-6(ev400)* (63% and 62%) (Figure 3H and I). These axon guidance data suggest that *unc-6(lq154)* might more strongly affect the long-range guidance of axons toward the ventral nerve cord.

**Figure 3.**
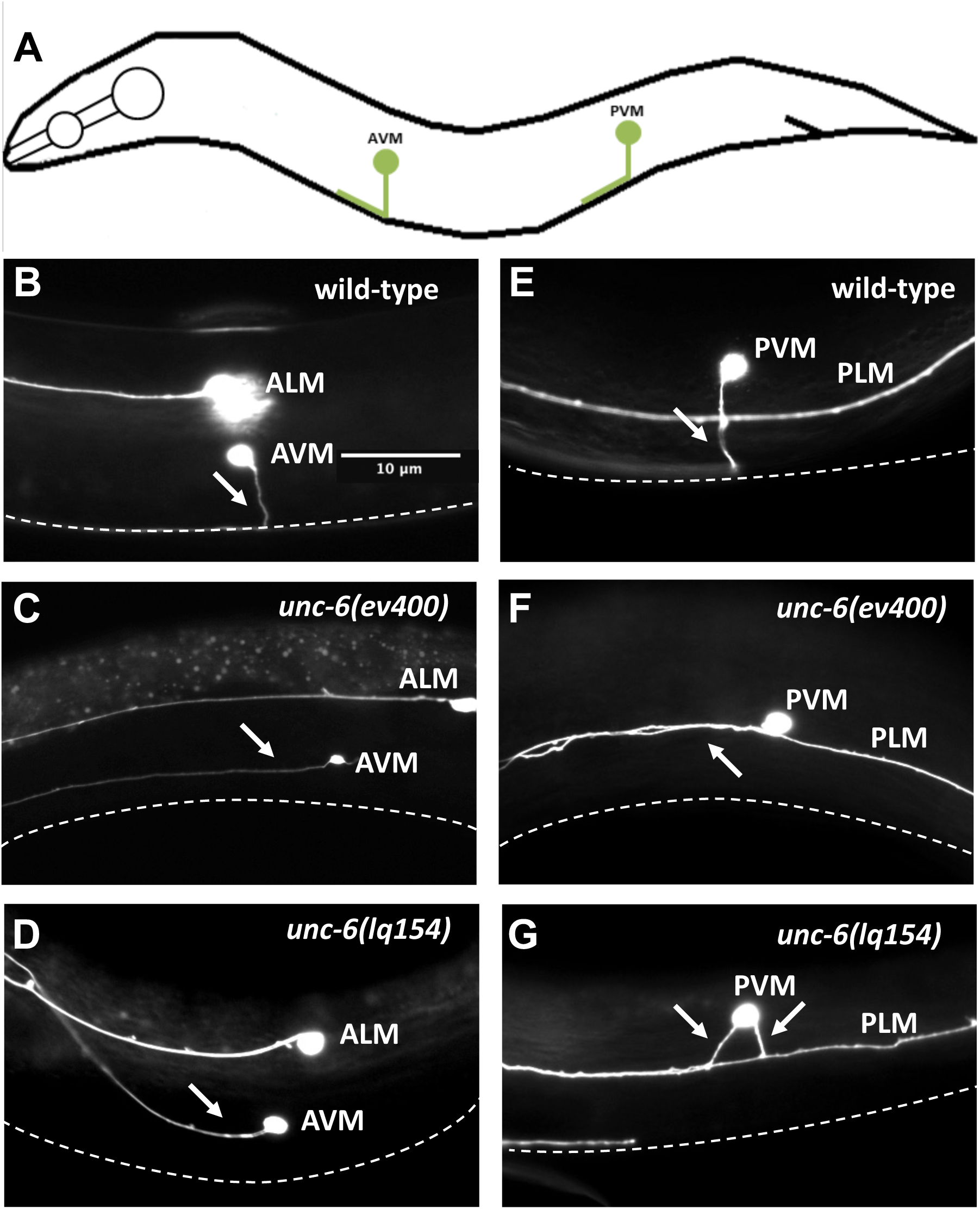
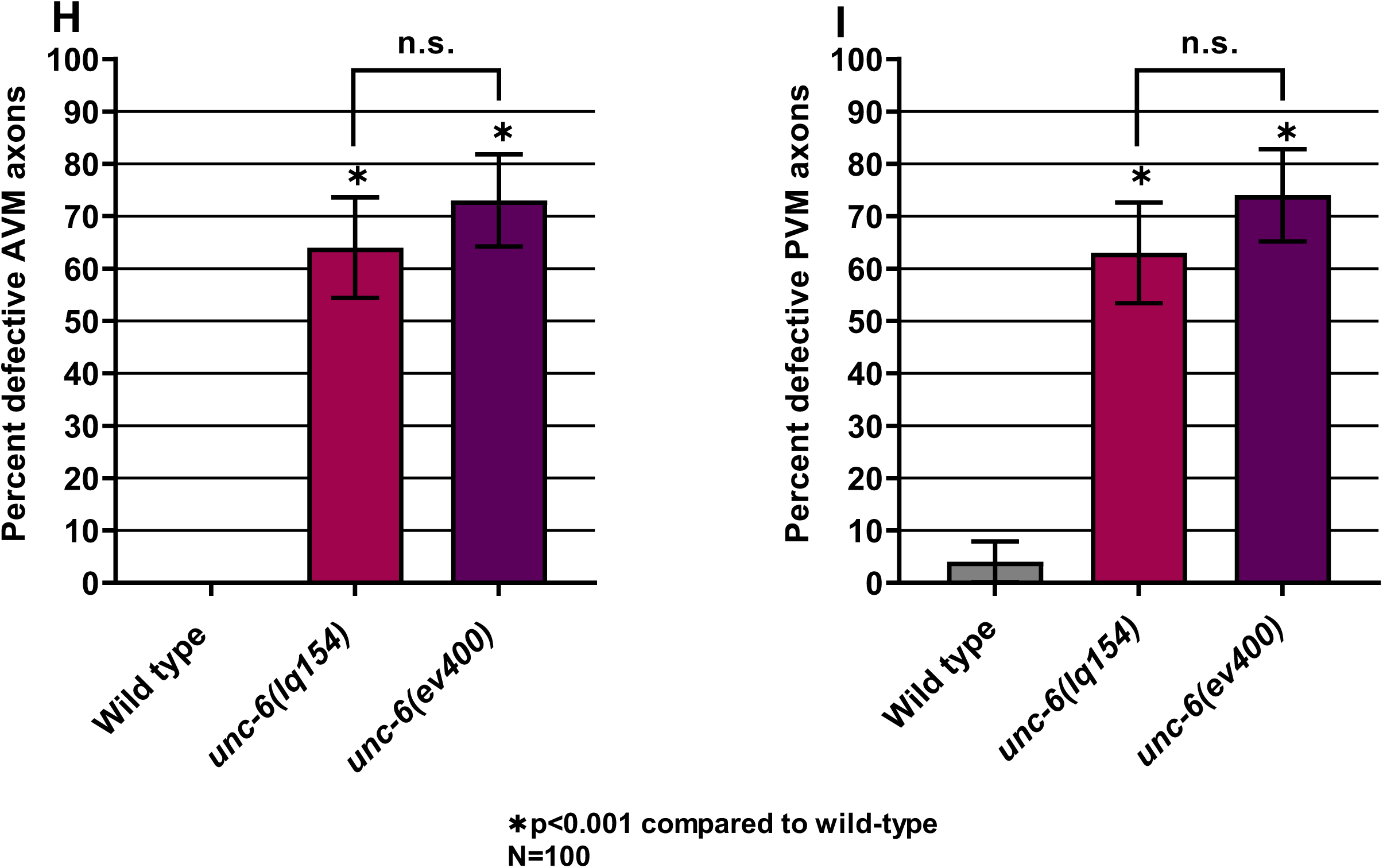
AVM/PVM axon guidance defects in *unc-6* mutants. A) A diagram illustrating AVM and PVM structure. Anterior is to the left and dorsal is up. The AVM and PVM cell bodies are located laterally and extend axons that migrate ventrally to join and run anteriorly within the ventral nerve cord. B-C) Fluorescent micrographs of AVM and PVM neurons expressing the *zdIs4[Pmec-4::gfp]* transgene in wild-type and *unc-6* mutants. The ALM and PLM cell bodies and processes are also indicated. The ventral nerve cord is indicated by a white dashed line. In *unc-6* mutants, AVM and PVM axons fail to extend ventrally and instead extend laterally, failing to reach the VNC. The scale bar in B represents 10 μm for all images. H-I). Graphs showing the percentage of defective AVM and PVM axons (failure to reach the ventral nerve cord, branching, or turning at >45° angle during migration) in *unc-6* mutants. Error bars represent 2x the standard error of proportion. Significance was determined using Fisher’s exact test.

### VD growth cones were initially polarized dorsally in *unc-6(lq154)*

The hypomorphic VD/DD dorsal axon guidance phenotype of *unc-6(lq154)* was investigated in more detail by analyzing VD growth cone polarity during their dorsal commissural outgrowth. VD growth cones in wild-type display a dorsal polarity of filopodial protrusions, such that ∼70% of filopodial protrusions occur from the dorsal half of the growth cone (Knobel *et al.* 1999; Norris and Lundquist 2011) (Figure 4A-C). This dorsal polarity of filopodial protrusion is abolished in *unc-6(ev400)* mutants, indicating that *unc-6* is required to polarize the VD growth cone during outgrowth (Norris and Lundquist 2011) (Figure 4B and F).

**Figure 4.**
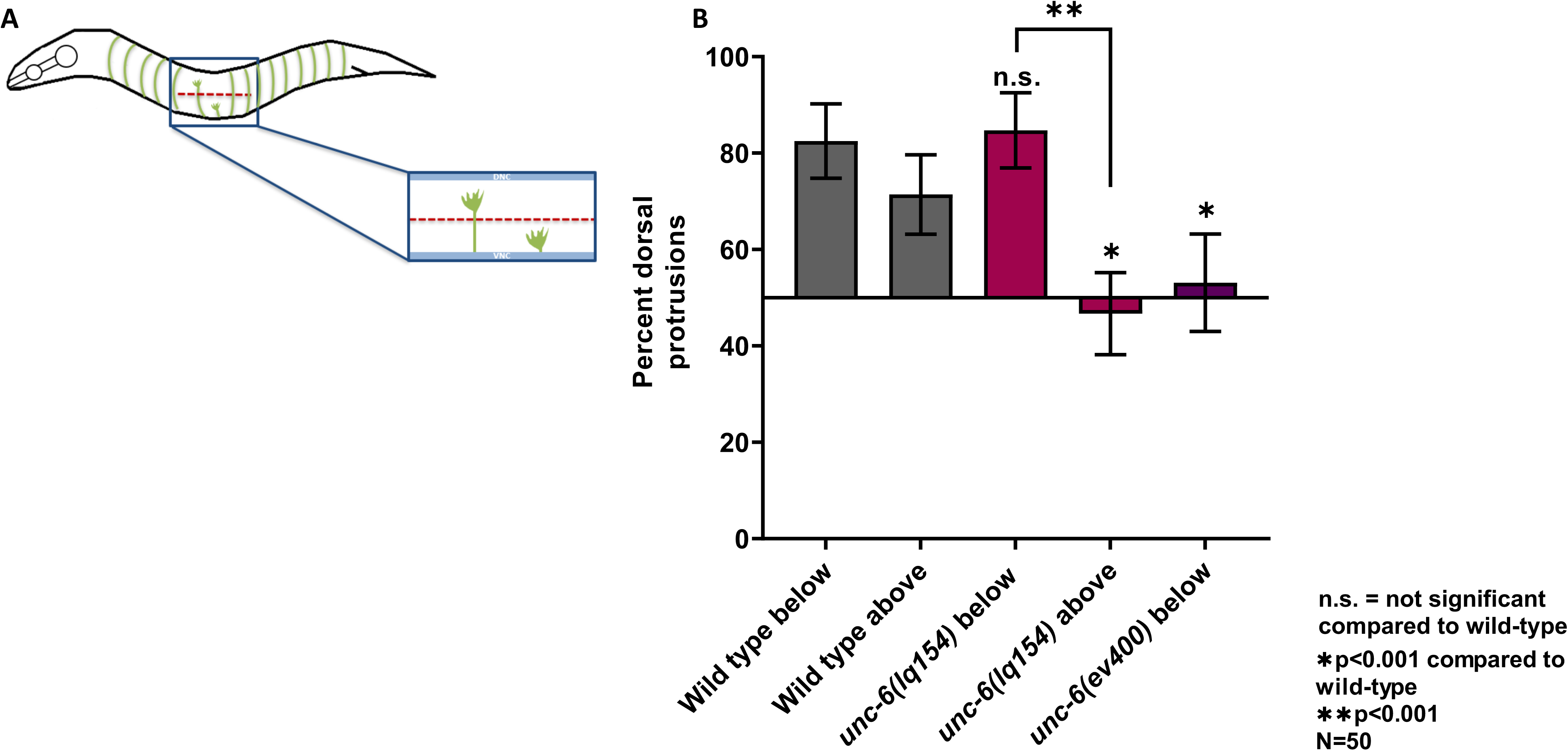
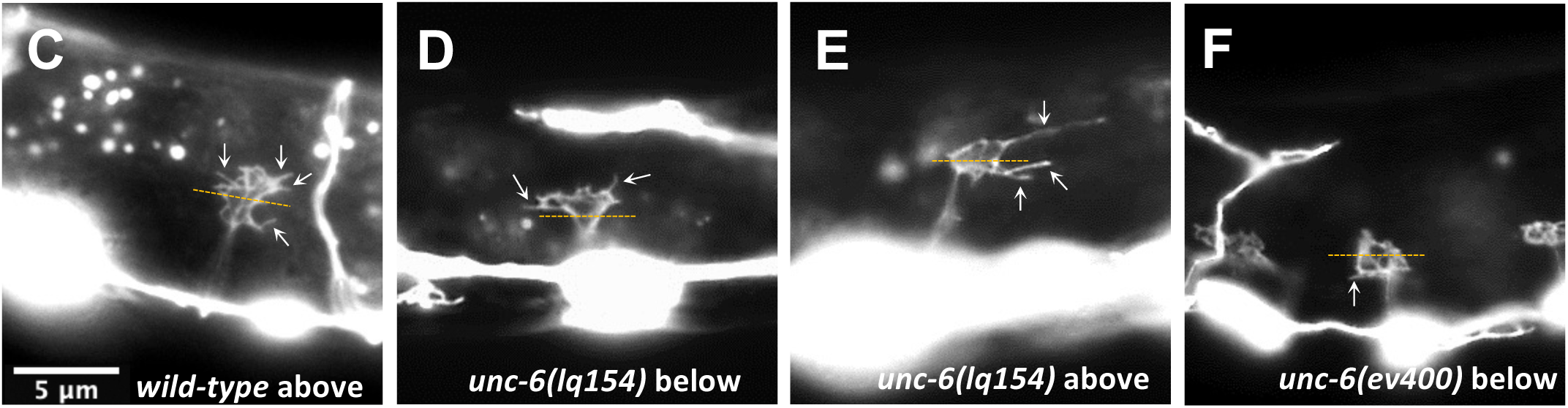
VD growth cone polarity in *unc-6* mutants. A) Diagram of early L2 *C. elegans* hermaphrodite illustrating VD/DD axons and VD growth cones. Anterior is to the left and dorsal is up. During early L2 VD/DD axons extend anteriorly in the ventral nerve cord before turning dorsally to extend to the dorsal nerve cord. The lateral midline is represented by the dashed red line and the thick blue lines represent the ventral and dorsal nerve cords. VD growth cones are shown both below and above the midline. (B) Percentage of dorsally biased filopodial protrusions of VD growth cones below and above the lateral midline. Significance between wild-type and mutants was determined by Fisher’s exact test. Error bars represent 2x the standard error of proportion. C-F) Representative images of VD growth cones expressing the *juIs76[Punc-25::gfp]* transgene. Polarity is determined by dividing the growth cone equally into dorsal and ventral subsections respective to the ventral nerve cord and counting the number of protrusions. The dashed yellow line represents the approximate midline used to divide the growth cone into equal sections. White arrows point to filopodia. The scale bar in C represents 5 μm for all images. C) Wild-type growth cones display dorsally biased filopodial protrusions. D-E) *unc-6(lq154)* does not affect polarity below the midline but disrupts polarity above the midline.

Significantly more VD/DD axons extended past the lateral midline in *unc-6(lq154)* compared to the *unc-6(ev400)* null, suggesting that guidance defects in *unc-6(lq154)* were occurring after lateral midline crossing. Therefore, growth cones at different points in their migration were analyzed, specifically those that were below the lateral midline and those that were above (Figure 4A). Wild-type VD growth cones were dorsally polarized both below and above the lateral midline (82% and 70%, respectively) (Figure 4B and C). In *unc-6(ev400)* null mutants, most VD growth cones did not extend past the lateral midline. Those below the lateral midline showed a loss of polarity (52%) (Figure 4B and D). In contrast, *unc-6(lq154)* VD growth cones were still significantly polarized below the lateral midline (84%) (Figure 4B and D). However, polarity was abolished in *unc-6(lq154)* VD growth cones above the midline (48%) (Figure 4B and E). These results suggest that in *unc-6(lq154)*, VD growth cones were initially polarized, but this polarity was not maintained as dorsal outgrowth proceeded. This could explain the hypomorphic nature of *unc-6(lq154)* in dorsal VD axon guidance.

F-actin accumulation is also dorsally-polarized in wild-type VD growth cones, which is abolished in *unc-6(ev400)* null mutants (Norris and Lundquist 2011) (Figure 5A-D). F-actin was visualized using the VAB-10 F-actin binding domain fused to GFP as previously described (Norris and Lundquist 2011). F-actin was dorsally polarized in *unc-6(lq154)* VD growth cones below the lateral midline (Figure 5A and E-G), but was unpolarized in those above the lateral midline (Figure 5A and H-J). These results further suggest that VD growth cones are initially polarized in *unc-6(lq154)*, and that polarity is lost as the growth cones migrate dorsally.

**Figure 5.**
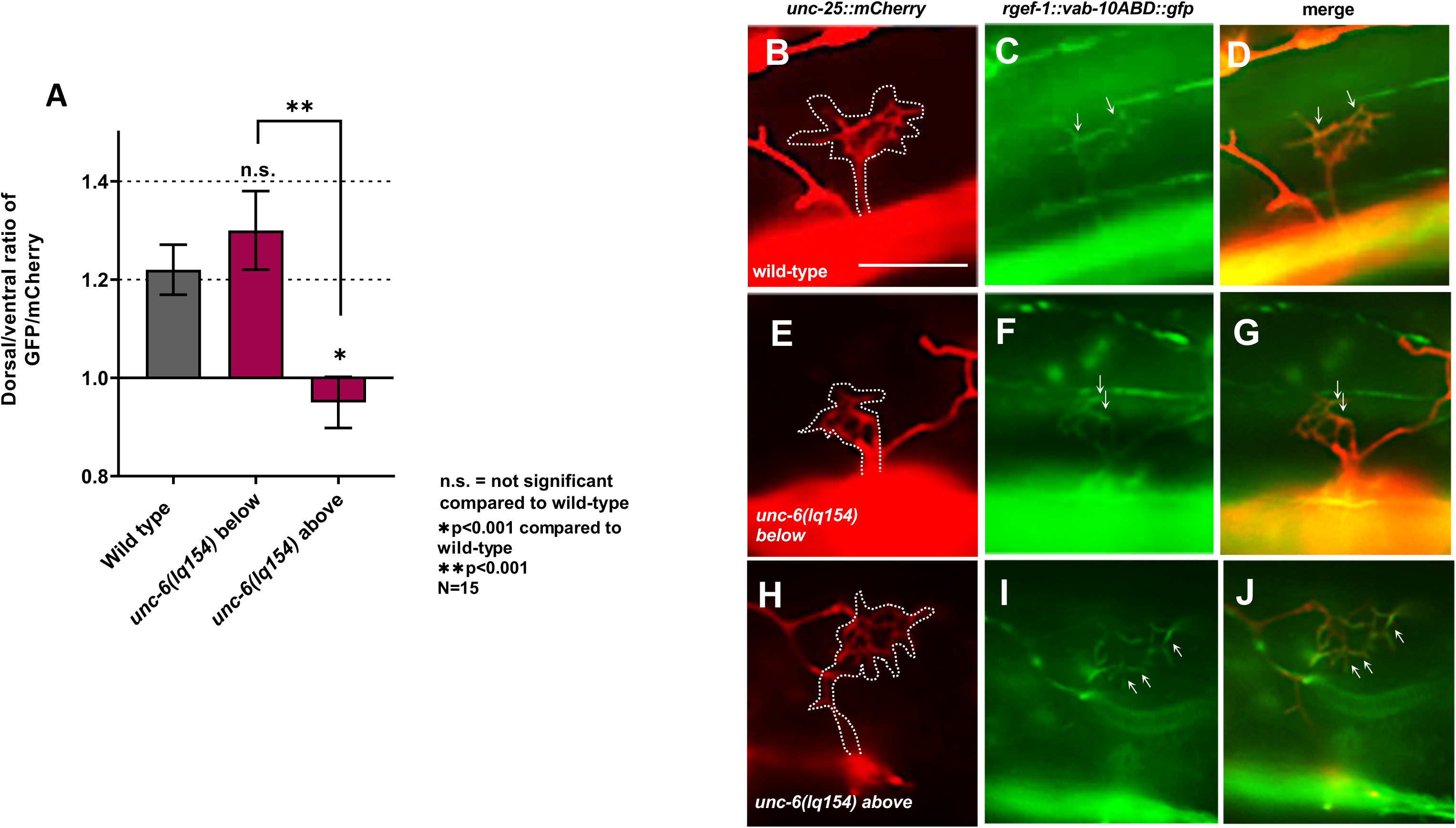
F-actin polarity is lost above the midline in *unc-6(lq154).* (A) Graph showing the dorsal-to-ventral ratio of GFP/mCherry in VD growth cones expressing an mCherry volume (*lhIs6[Punc-25::mCherry]* marker and the VAB-10 F-actin binding domain fused to GFP (*lqIs170* [*Prgef-1::vab-10ABD::gfp*]). Five line scans from dorsal to ventral were drawn across each growth cone, and the intensities of mCherry and GFP were determined in dorsal and ventral sections. Error bars represent standard error of the mean. Statistical significance was determined using a two-tailed *t*-test with unequal variance. B-J) Images of VD growth cones with mCherry volume marker, VAB-10ABD::GFP, and a merge. The dotted line in the mCherry micrographs represents the approximate area of the growth cone. The scale bar in B represents 5 μm for all images. B-D) A wild-type VD growth cone. Arrows point to dorsally-localized GFP. E-G) An *unc-6(lq154)* growth cone below the lateral midline. Arrows point to dorsally-localized GFP. H-J) An *unc-6(lq154)* growth cone above the lateral midline. Arrows point to ventrally located GFP indicating a loss of dorsal polarity of F-actin.

### Anterior and dorsal *unc-6* ectopic expression rescued VD/DD axon guidance defects and VD growth cone polarity in *unc-6* mutants

At the time of VD outgrowth, *unc-6* is expressed in the VA and VB motor neurons in the ventral nerve cord, which could be the source of UNC-6 that directs the VD growth cones dorsally. In electron microscopic reconstruction of the ventral nerve cord (White *et al.* 1986), the VA and VB neurons and processes are often ventral to and in contact with the VD/DD neurons and processes in the ventral nerve cord (Figure 6A).

**Figure 6.**
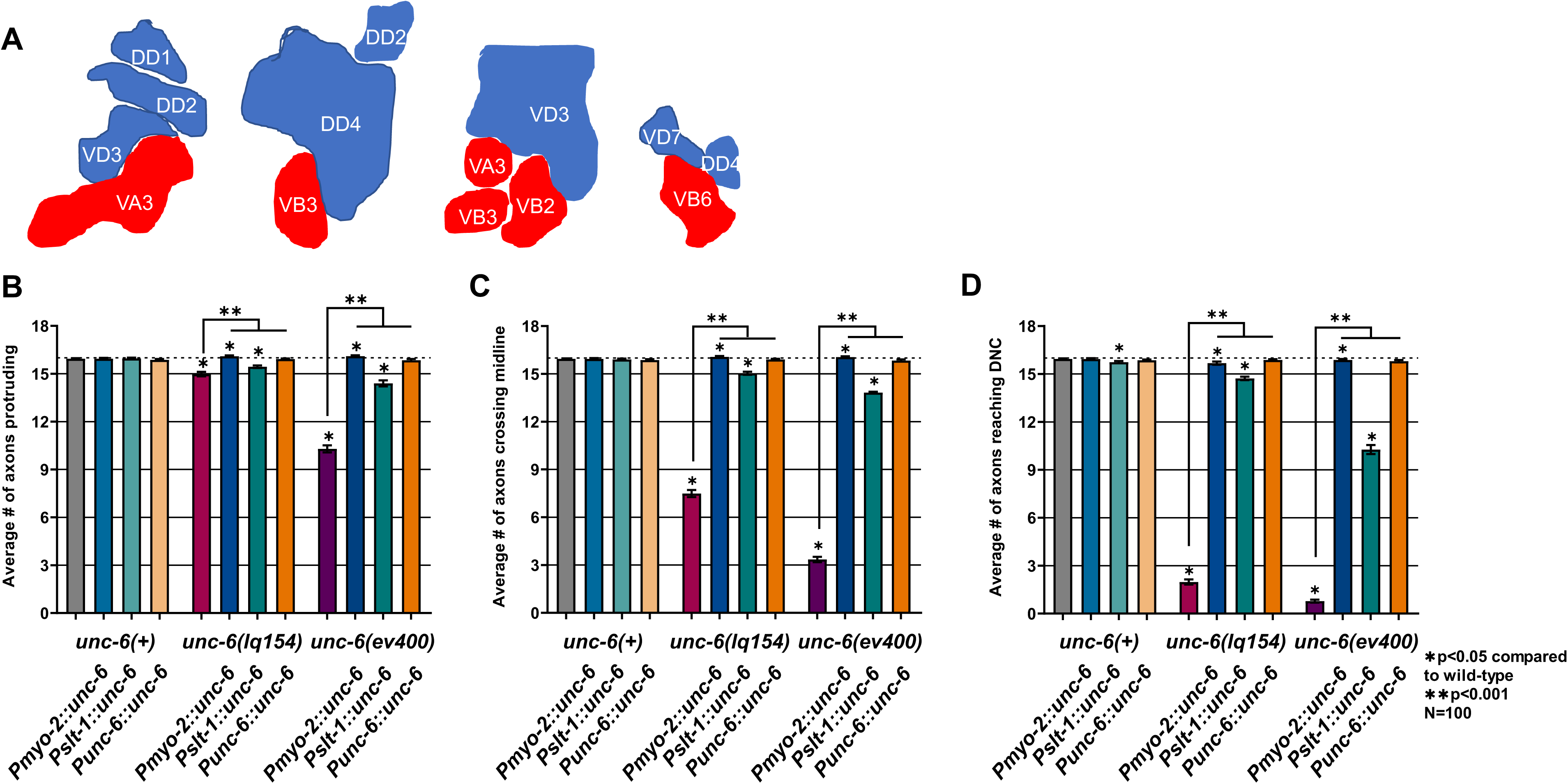
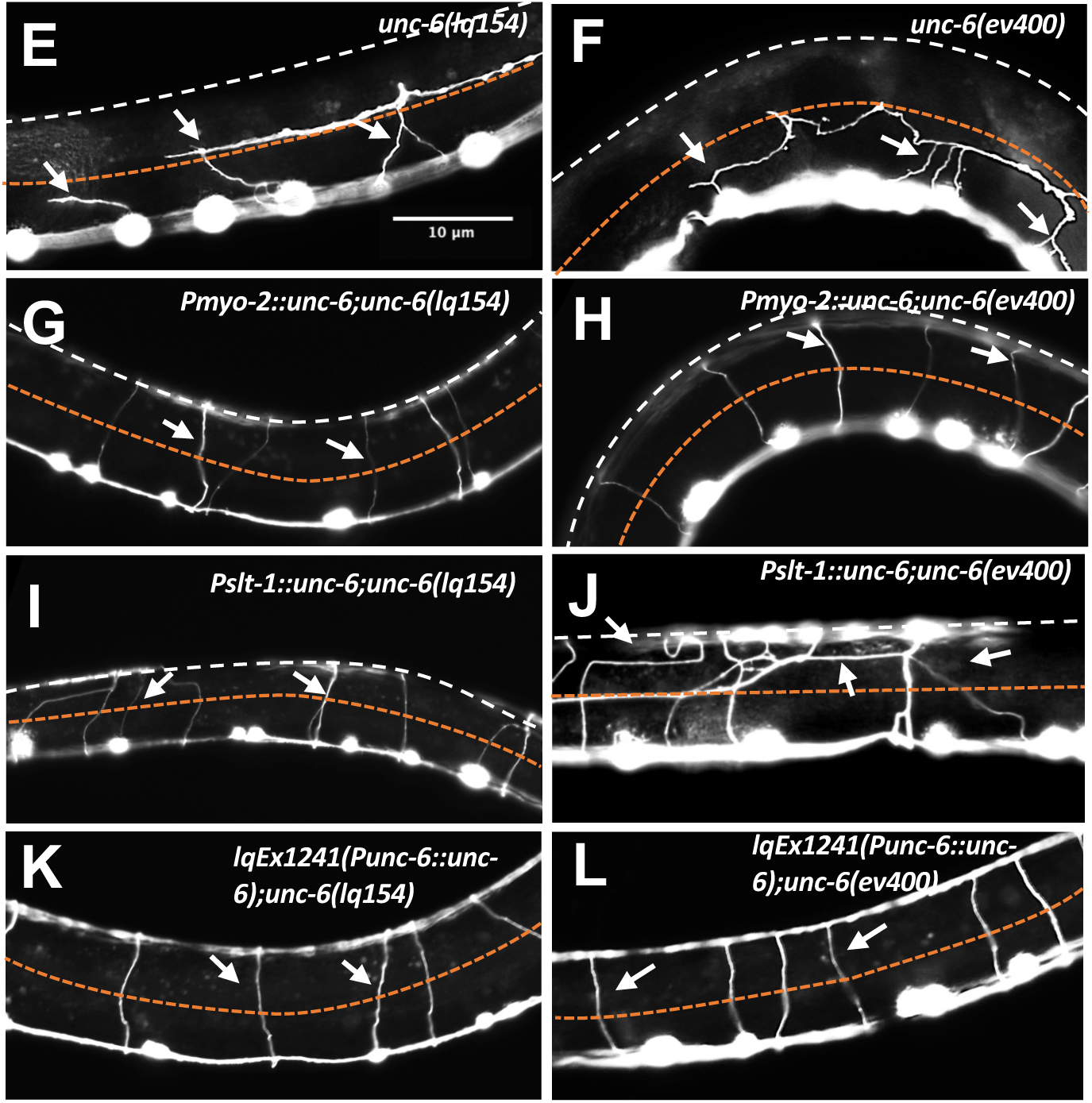
*unc-6* ectopic expression rescues VD/DD axon guidance defects of *unc-6(lq154)* and *unc-6(ev400)* mutants. A) The figures are tracings of transmission electron microscopic sections of the ventral nerve cord showing the relative positions of the VA and VB neurons expressing *unc-6* (red) and the VD/DD neurons (blue). Dorsal is up. The image tracings were derived from White et al., 1986, *The Structure of the Nervous System of the Nematode* Caenorhabditis elegans. From left to right: p. 180, panel D; p. 338, panel A; p. 331, panel D; and p. 335, panel C. B) Graphs showing the average number of axons protruding from the ventral nerve cord, passing the lateral midline, or reaching the dorsal nerve cord, as described in Figure 2. Error bars represent standard error of the mean. Statistical significance was determined by a two-tailed *t*-test with unequal variance. E-L) Representative images of VD/DD axons in indicated genotypes. Approximate lateral midline is indicated by the dashed orange line and the dorsal nerve cord is indicated by the dashed white line. White arrows point to axons. Anterior is to the left and dorsal is up. The scale bar in B represents 10 μm for all images.

To test if a ventral source of UNC-6 was required for dorsal VD/DD axon pathfinding and VD growth cone polarity, transgenes that express wild-type *unc-6(+)* from ectopic, non-ventral sources were constructed. A *Pmyo-2::unc-6(+)* transgene was predicted to drive UNC-6 expression in the pharyngeal muscle in the anterior of the animal (Ardizzi and Epstein 1987; Fire *et al.* 1990); and a *Pslt-1::unc-6(+)* transgene was predicted to express UNC-6 in dorsal body wall muscle (Hao *et al.* 2001).

In a wild-type *unc-6(+)* background, *Pmyo-2::unc-6* and *Pslt-1::unc-6* expression caused no significant defects in VD/DD axon guidance and had no significant effect on VD growth cone polarity (Figure 6B-D and Figure 7A and B), suggesting that these transgenes were not significantly interfering with endogenous *unc-6* function. Surprisingly, *Pmyo-2::unc-6* and *Pslt-1::unc-6* expression strongly rescued VD/DD axon guidance defects and uncoordinated locomotion of both *unc-6(ev400)* null mutants and *unc-6(lq154)* mutants (Figure 6B-D and E-L). VD growth cone dorsal polarity of filopodial protrusion was also restored in both *unc-6(ev400)* and *unc-6(lq154)* mutants (Figure 7). Rescue by *Pmyo-2::unc-6* and *Pslt-1::unc-6* did not differ significantly from rescue of *unc-*6 driven by the endogenous *unc-6* promoter (*Punc-6::unc-6*) (Figure 6B-D and Figure 7A and B). These results indicate that ectopic expression of *unc-*6 from non-ventral sources can rescue VD/DD dorsal axon guidance and VD growth cone dorsal polarity of filopodial protrusion in *unc-6* mutants and that a strictly ventral source of UNC-6 is not required.

**Figure 7.**
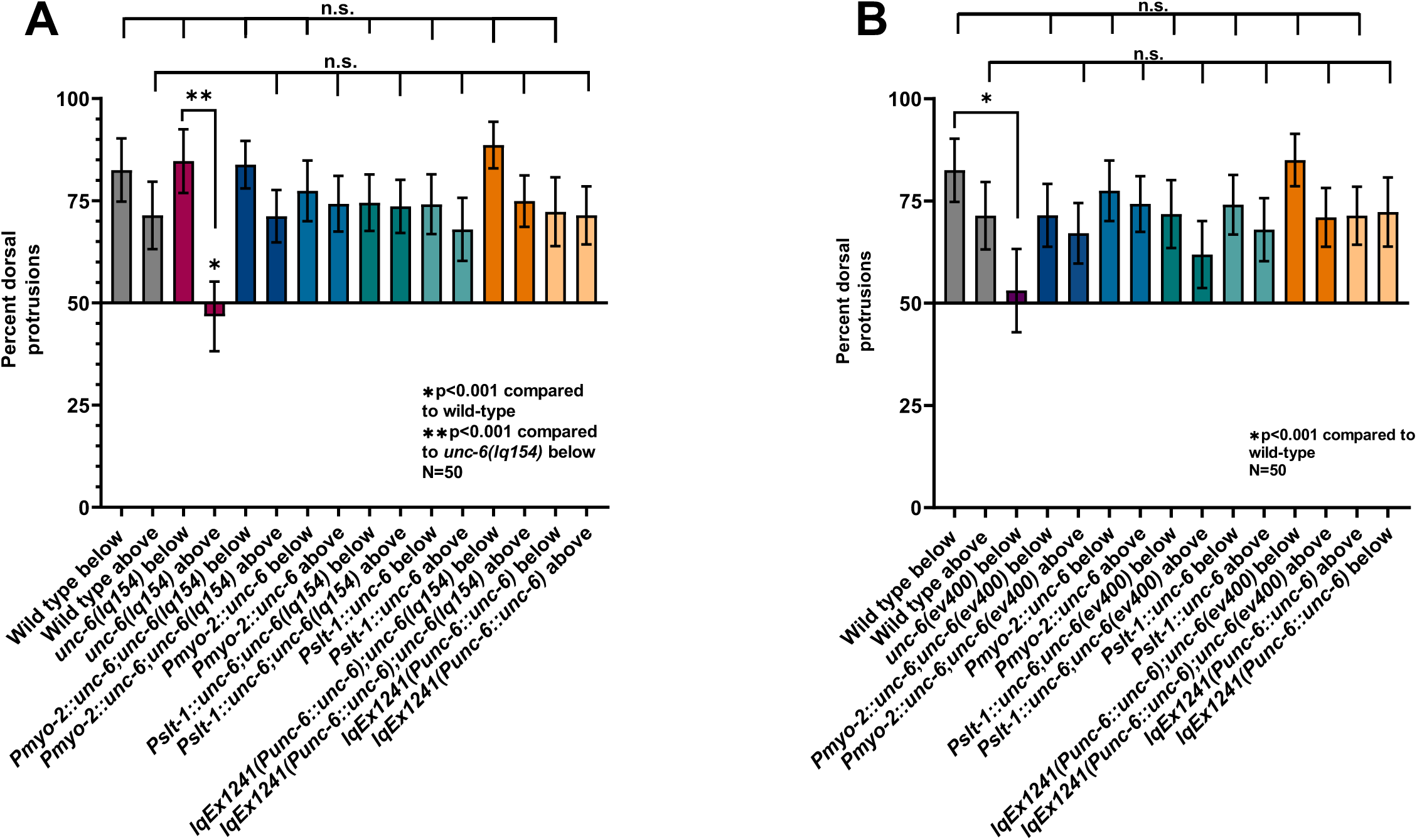
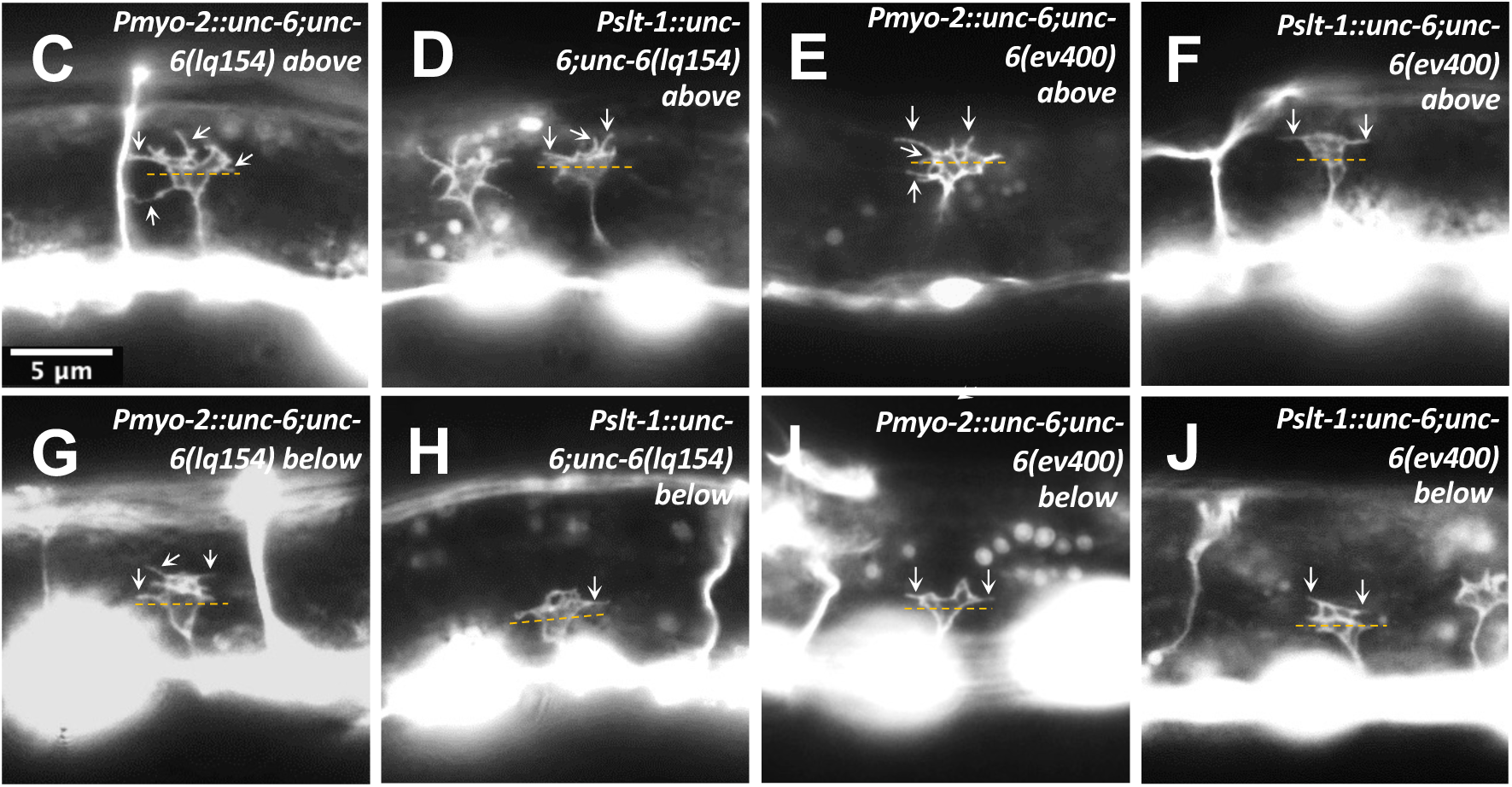
VD growth cone polarity defects of *unc-6* mutants are rescued by *unc-6* ectopic expression. A-B) Graphs of VD growth cone polarity as described in Figure 4. Error bars represent 2x the standard error of proportion. Statistical significance was determined using Fisher’s exact test. C-J) Fluorescent micrographs of VD growth cones. Arrows point to filopodial protrusions. Approximate midline used to divide the growth cone into equal sections is indicated by dashed yellow line. The scale bar in C represents 5 μm for all images.

### AVM/PVM ventral axon guidance does not require a ventral UNC-6 source

The above results suggest that dorsally-directed axon guidance of VD/DD neurons does not require a ventral UNC-6 source. The AVM and PVM axons grow ventrally and also require UNC-6 for their ventral guidance (Figure 3).

*Pmyo-2::unc-6* and *Pslt-1::unc-6* expression significantly rescued AVM/PVM ventral axon guidance in both *unc-6(ev400)* null and *unc-6(lq154)* (Figure 8). These results indicate that a ventral source of UNC-6 is not required for ventral axon guidance. Together with the VD/DD results, these data indicate that an asymmetric ventral source of UNC-6 is not required for dorsal-ventral axon guidance.

**Figure 8.**
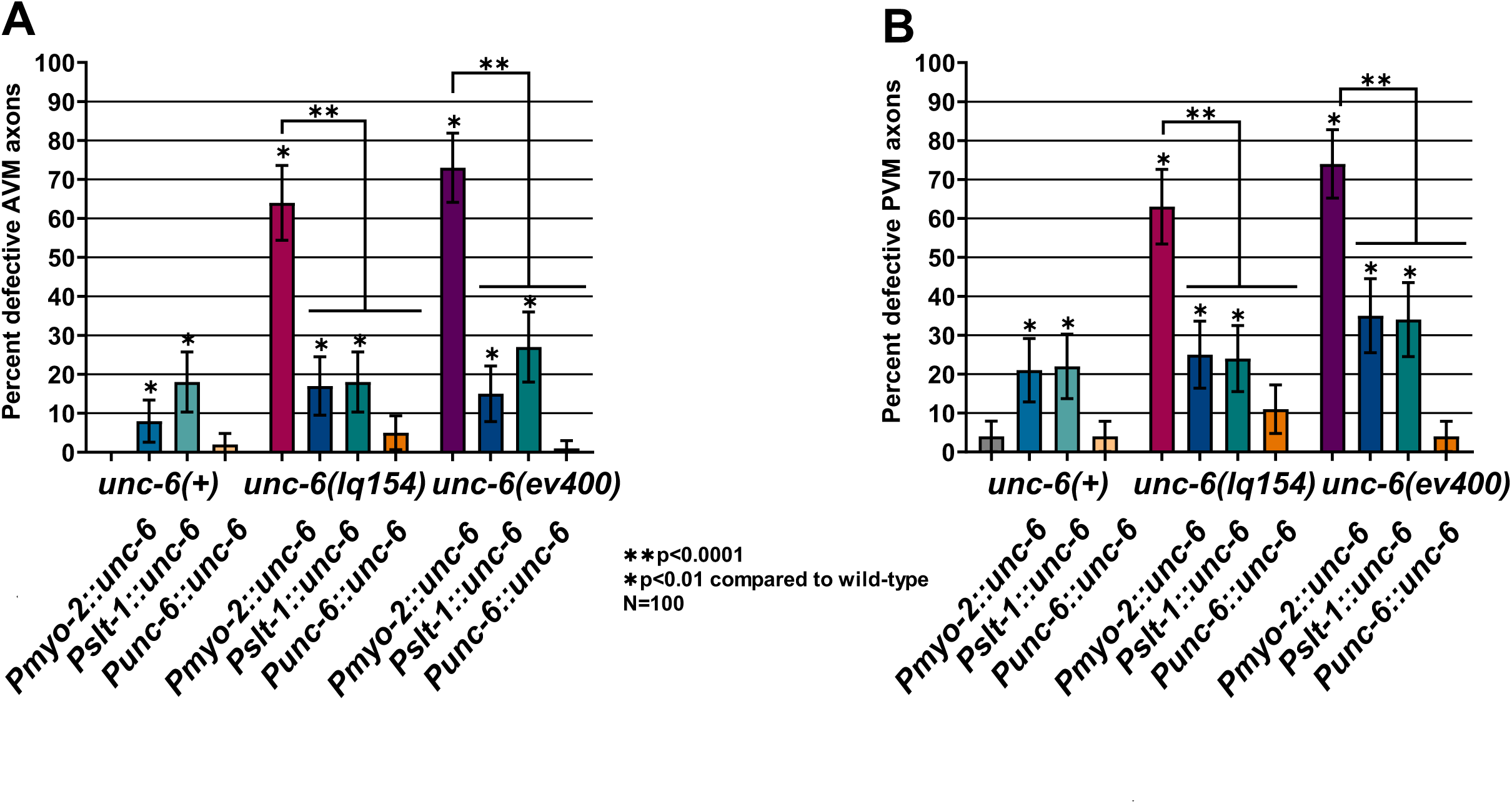
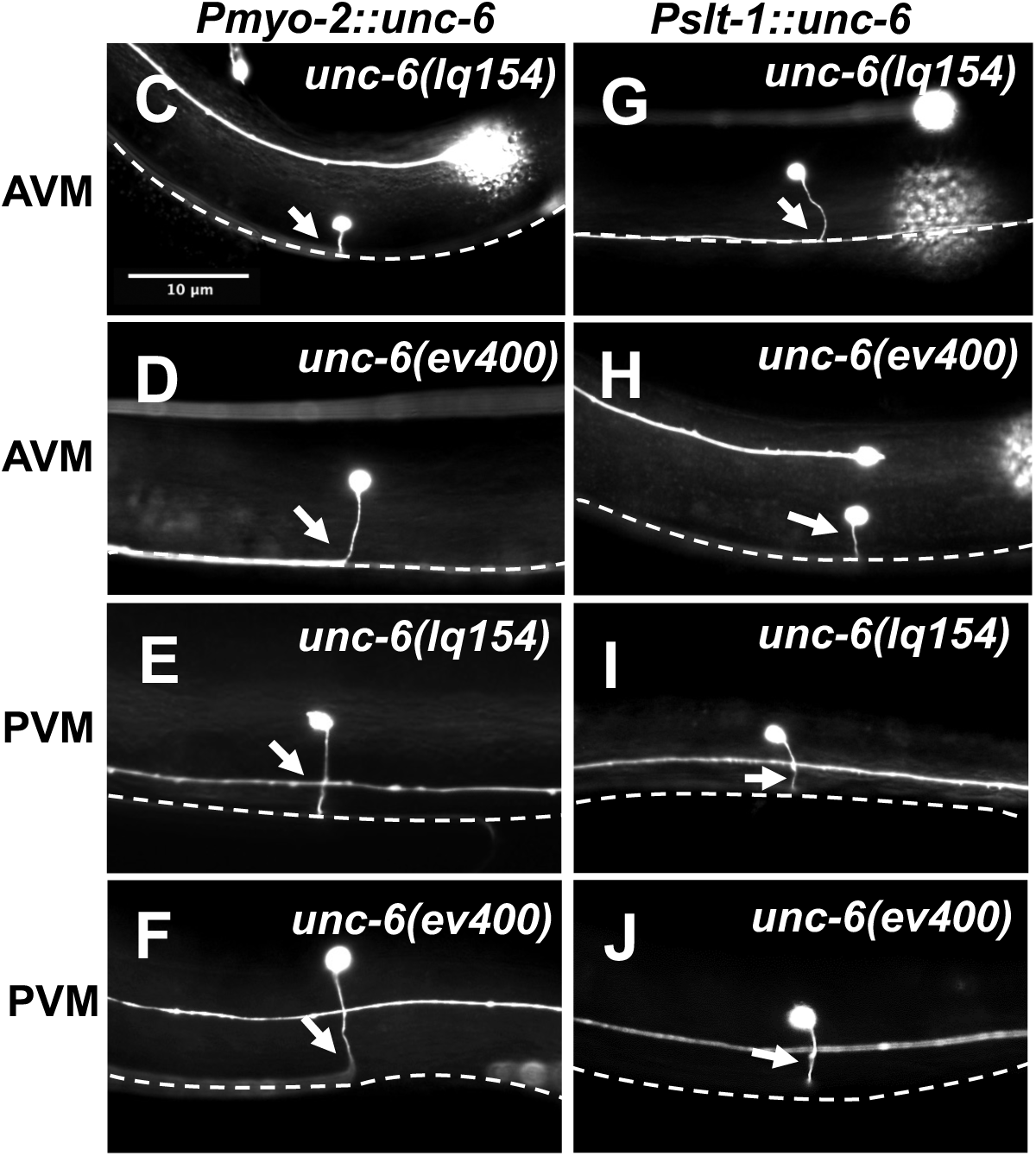
*unc-6* ectopic expression rescues AVM/PVM axon guidance defects of *unc-6* mutants. (A and B) Graphs showing the percentage of defective AVM and PVM axons (failure to reach ventral nerve cord, branching, or turning at >45° angle during migration) as described in Figure 3. Error bars represent 2x the standard error of proportion. Statistical significance was determined by Fisher’s exact test. C-J) Representative images of AVM and PVM neurons as indicated. White arrows indicate axons. C-F) *unc-6* mutants with the *Pmyo-2::unc-*6 transgene. G-J *unc-6* mutants with the *Pslt-1::unc-6* transgene. The ventral nerve cord is indicated by dashed white line. The scale bar in C represents 10 μm for all images.

In a wild-type background, *Pmyo-2::unc-6* and *Pslt-1::unc-6* caused significant defects in AVM and PVM ventral axon guidance (8-22%) (Figure 8). Expression from the endogenous *unc-6* promoter did not perturb AVM/PVM ventral guidance (Figure 8). This suggests that ectopic expression of *unc-6* might interfere with the ventral guidance of AVM and PVM axons in a wild-type background.

### HSN ventral axon guidance does not require a ventral UNC-6 source

The axon of the HSN neuron required *unc-6* for ventral growth (Figure 9). In the SOAL model of HSN axon guidance, the UNC-40 receptor forms spontaneous membrane clusters that are oriented to the ventral surface of the HSN cell body by UNC-6, defining the site of HSN axon formation (Kulkarni *et al.* 2013; Yang *et al.* 2014; Limerick *et al.* 2017). Both *unc-6* null and *unc-6(tm)* mutants show 87% HSN ventral axon guidance defects, where the HSN axon migrates laterally and fails to migrate ventrally (Figure 9). *Pmyo-2::unc-6* and *Pslt-1::unc-6* strongly rescued the HSN axon guidance defects of both mutants (Figure 9A). This indicates that a strictly ventral source of *unc-6* is not required for HSN ventral guidance and suggests that the SOAL model does not require strictly ventral UNC-6.

**Figure 9.**
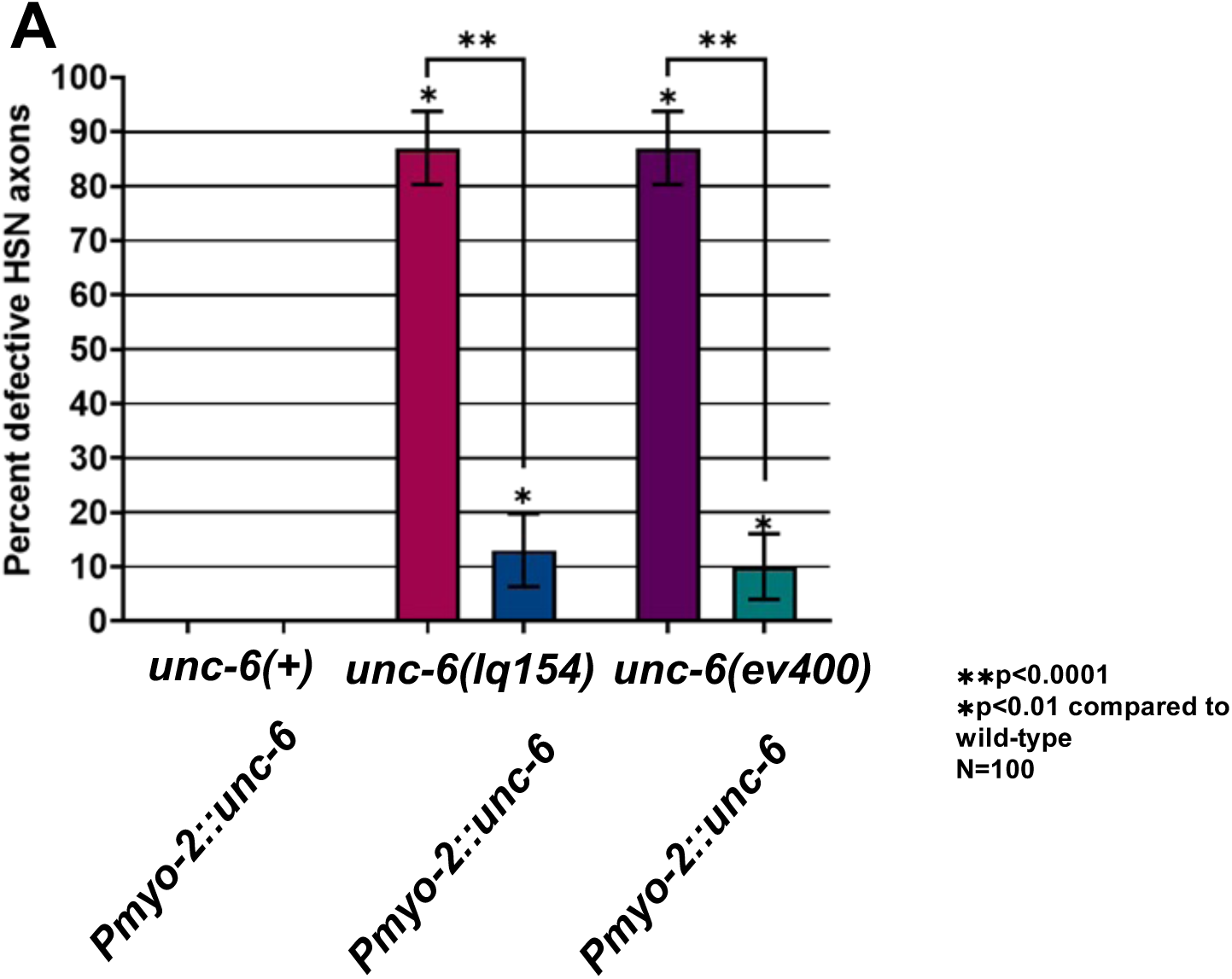
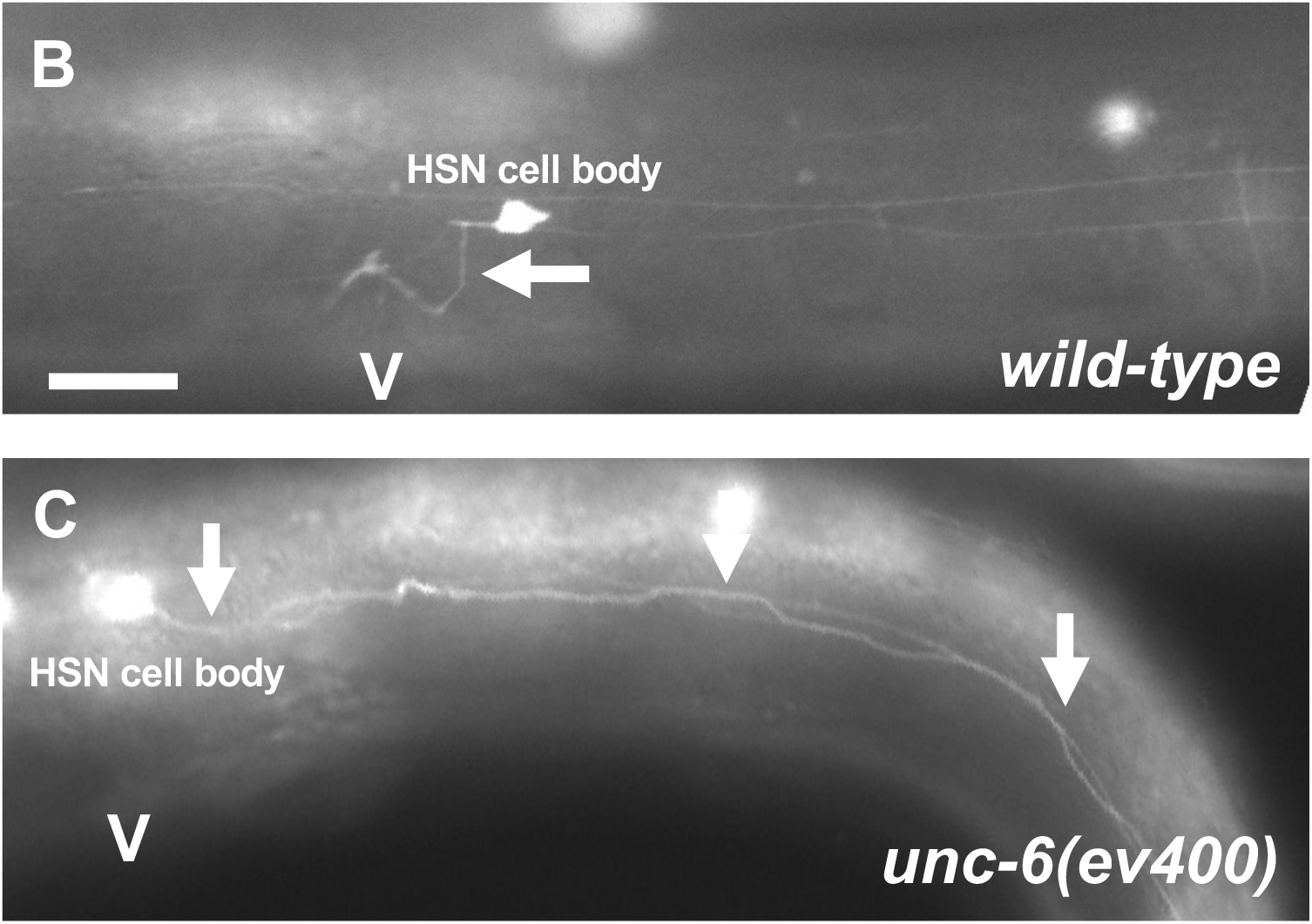
Ectopic *unc-6* rescues HSN ventral guidance defects. A) A graph of the percent of HSN ventral guidance defects in given genotypes. B and C) Fluorescent micrographs of HSN axons expressing *kyIs179[Punc-86::gfp].* The scale bar in B represents 5 μm. A “V” indicates the position of the vulva. B) In wild-type, the HSN axon (arrow) extends anteriorly before turning ventrally toward the vulva. C) In *unc-6(ev400)* an HSN axon (arrows) extends posteriorly and fails to turn ventrally.

**Figure 10.**
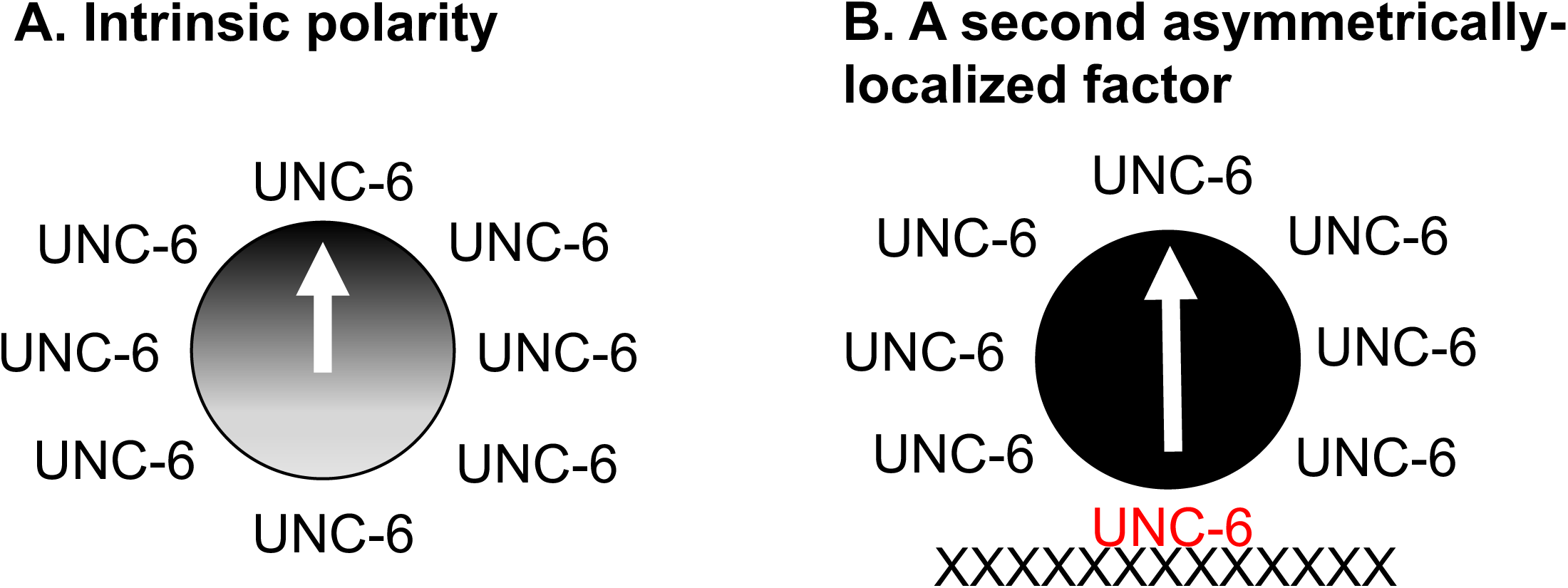
Models of ectopic UNC-6 in dorsal-ventral axon guidance. A) The cell has intrinsic polarity and UNC-6 acts as a permissive cue. B) To polarize the cell, UNC-6 must interact with a second, unidentified factor that is asymmetrically distributed (Xs).

## Materials and Methods

### Genetic methods

Experiments were performed at 20°C using standard *C. elegans* techniques (Brenner, 1974). Mutations used were *lqEx1241* [*Punc-6::Venus::unc-*6]; LGII: *juIs76* [*Punc-25::gfp*]; LGIV: *zdIs4* [*mec-4::gfp*]; LG V: *kyIs179 [Punc-86::gfp].* LGX: *unc-6(ev400, lq154), lqIs170* [*Prgef-1::vab-10ABD::gfp*], *lqIs379* [*Pmyo-2::Venus::unc-6*], *lhIs11* [*Pacr-2::mCherry*]. Chromosomal locations not determined: *lqIs401* [*Pslt-1::Venus::unc-6*] and *lhIs6* [*Punc-25::mCherry*]. Transgenic strains were obtained by microinjection of plasmid DNA into the germline. Multiple extrachromosomal transgenic lines of *Pmyo-1::Venus::unc-6* and *Pslt-1::Venus::unc-6* were established and integrated into the genome via standard techniques (Mellow and Fire, 1995). One integrated strain of each was chosen for further analysis. *lqEx1241* [*Punc-6::Venus::unc-6*] was maintained as an extrachromosomal array. Mutations were confirmed using PCR genotyping and sequencing. Wormbase (Davis *et al.* 2022) was utilized for *C. elegans* informatics.

### Transgene construction

Transgenes were created using a *pVns::unc-6* plasmid containing the *unc-6* coding region as well as the upstream promoter and 3’ UTR regions (Asakura et al., 2007). The *unc-6* promoter was replaced in with the *myo-*2 and *slt-1* promoters. Each plasmid was confirmed using PCR and sequencing.

### CRISPR/Cas9 to generate *unc-6(lq154)*

CRISPR/Cas9 genome editing was used to insert a transmembrane signal sequence into the 3’ end of *unc-6.* sgRNAs were engineered to target the 3’ end of exon 12 and the 3’ UTR. A donor plasmid containing left and right homology arms with recoded sgRNA regions, GFP, a synthetic intron flocked by LoxP sites that contains a hygromycin resistance cassette, a re-coded exon 13, and the coding region for the *cdh-3* transmembrane segment was used as a repair template. The repair plasmid sequence is the sequence of the genome edit (pnu1884; Supplemental File 1). A mixture of sgRNAs, Cas9, and donor homology plasmid was injected into the gonads of N2 animals. Plates were treated with hygromycin and surviving animals were screened for the insert by PCR genotyping. The *unc-6(lq154)* genome edit was confirmed by sequencing the region from *unc-6(lq154)* genomic DNA. Genome editing reagents were provided by InVivo Biosystems (Eugene, OR, USA). sgRNA 1: ATTATGGATAAGgtaagaac sgRNA 2: agattggatcaggagtcaca

### RNA seq on unc-6(lq154)

Total RNA was isolated from mixed stage *unc-6(lq154)* animals (strain LE5443, *unc-6(lq154) X; juIs76 II*) as described previously. RNA seq libraries were constructed using the NEBNext stranded mRNA library kit. Sequencing was conducted on a Nextseq 2000 instrument with paired-end 150-bp sequencing. Preprocessing of FASTQ files was completed using fastp (0.23.2) (Chen *et al.* 2018), which included adapter trimming, per-read cutting by quality score, global trimming, and filtering out bad reads. HISAT2 (version 2.2.1) (Kim *et al.* 2015) was used to align reads to a *C. elegans* reference genome [release WBcel235, version WBPS14 (WS271)] that had been edited to include the *unc-6(lq154)* genome edit sequence. Resulting BAM files were analyzed in the Integrated Genome Viewer (Robinson *et al.* 2011; Thorvaldsdottir *et al.* 2012). Raw reads can be found in the Sequence Read Archive PRJNA1093133.

### Imaging and quantification of axon guidance defects

The AVM and PVM axons were visualized with a *Pmec-4::gfp* transgene, *zdIs4,* which is expressed in the touch receptor neurons (Royal et al., 2005). AVM and PVM axons were considered defective if they failed to reach the ventral nerve cord, wandered laterally at more than a 45° angle during ventral migration, or had ectopic processes. HSN axons were scored using the *Punc-86::*gfp transgene *kyIs179.* HSN axons were considered defective if they failed to extend ventrally to the ventral nerve cord. Axons were scored in L4 or pre-gravid adults. 100 animals were scored for each genotype and the percent defective axons was determined. Significance was determined by using a Fisher’s exact test. VD/DD neurons were visualized using a *Punc-25::gfp* transgene, *juIs76,* which is expressed in all GABAergic neurons (Jin et al., 1999). Of the 19 VD/DD axons, 18 extend commissures on the right side of the animal. The VD1 commissure on the left side was not scored. In wild-type animals, an average of 16 axons are observed due to fasciculation. Three reference points were used to quantify axon guidance defects: protrusion from the ventral nerve cord, crossing the lateral midline, and reaching the DNC. In mutants where less than 16 axons were observed only the observable axons were scored. In mutants where more than 16 axons were observed all observable axons were scored. Significance between genotypes was determined using a two-tailed t-test with unequal variance. 100 animals were scored per genotype.

DA/DB neurons were visualized using a *Pacr-2::mCherry* transgene, *lhIs11,* which is expressed in cholinergic motor neurons (provided by Dr. Brian Ackley). Of the 16 DA/DB axons, three are obscured by the *lhIs11* pharyngeal marker so they were not scored. An average of 12-13 axons are observed in wild-type animals due to fasciculation. Three reference points were used to quantify axon guidance defects: protrusion from the ventral nerve cord, crossing the lateral midline, and reaching the DNC. In all mutants, only observable axons were scored. Significance between genotypes was determined using a two-tailed t-test with unequal variance. 100 animals were scored per genotype.

### Growth cone imaging and quantification

VD growth cones were visualized using a *Punc-25::gfp* transgene, *juIs76* (Jin et al., 1999). Growth cones were imaged as previously described (Norris and Lundquist, 2011). Animals were collected at approximately 16 hours post-hatching and placed on a 2% agarose pad with 5mM sodium azide in M9 buffer. Mutants were harvested at different timepoints to account for differences in development time. Growth cones were imaged using a Qimagine Rolera mGi camera on a Leica DM5500 microscope and analyzed with ImageJ. To determine growth cone polarity, the growth cone was divided in half into dorsal and ventral subsections relative to the ventral nerve cord. Percent dorsal protrusions was determined by counting the filopodial on the dorsal half and dividing by the total filopodia. Per genotype, 50 growth cones were analyzed below the midline and above the midline. Statistical significance was determined using a Fisher’s exact test.

### VAB-10ABD::GFP imaging

F-actin was analyzed as previously described (Norris and Lundquist, 2011). The F-actin binding domain of VAB-10 fused to GFP was used to visualize F-actin within the growth cone (Patel et al., 2008). To control for growth cone size and shape the growth cone was visualized using a soluble mCherry marker. F-actin accumulation was analyzed using ImageJ. For each growth cone, five line scans were drawn from the dorsal edge to the ventral (see results). Pixel intensity was measured as a function of distance from the dorsal edge. Pixel intensity was averaged and normalized to the volumetric mCherry fluorescence in line scans from the dorsal and ventral half of the growth cone. This normalized ratio was determined for 50 growth cones below and above the midline in *unc-6(lq154)* and 50 in wild-type. Statistical significance was determined using a two-tailed t-test with unequal variance on the average normalized ratios.

## Discussion

Previous studies have shown that UNC-6/Netrin directs dorsal VD growth cone outgrowth by first polarizing the growth cone via the UNC-5 receptor such that filopodial protrusions and F-actin are biased dorsally on the growth cone. UNC-6/Netrin then maintains this polarity as the growth cone migrates dorsally, with the UNC-40 receptor driving protrusion dorsally, and the UNC-5 receptor inhibiting protrusion ventrally, resulting in net dorsal outgrowth (Norris and Lundquist 2011; Norris *et al.* 2014; Gujar *et al.* 2017; Gujar *et al.* 2018; Gujar *et al.* 2019). The initial polarization of the growth cone by UNC-6/Netrin could involve a close-range interaction, whereas diffusible UNC-6/Netrin might be required for the longer-range maintenance of polarity. To test this model, an *unc-6(lq154)* genome edit was created that has the potential to encode a transmembrane version of UNC-6, which is not expected to diffuse and is expected to act only at short range.

### Short and long-range roles of UNC-6

*unc-6(tm)* behaved as a hypomorphic *unc-6* mutation, and was less severely uncoordinated and displayed fewer VD/DD commissural axon outgrowth defects compared to the *unc-6(null)*. VD growth cones near the ventral nerve cord (ventral to the lateral midline) were polarized similar to wild-type, with filopodial protrusions and F-actin biased dorsal-ward, whereas as *unc-6(null)* mutants were unpolarized at this point, and in fact very few extended dorsally past the lateral midline. VD growth cones further away from the ventral nerve cord, dorsal of the lateral midline, were unpolarized in *unc-6(tm)*. This suggests that *unc-6(tm)* growth cones were initially polarized normally, but lost polarity as they moved away from the ventral nerve cord, explaining the hypomorphic VD/DD axon guidance defects of *unc-6(tm)* mutants.

At the time of VD growth cone outgrowth, UNC-6 is expressed in the VA and VB motor neurons that extend axons in the ventral nerve cord (Wadsworth *et al.* 1996). In the adult ventral nerve cord, VD and DD axons are often in contact with and dorsal to the VA and VB axons (White *et al.* 1986). An intriguing hypothesis is that *unc-6(tm)* retains close-range functions of UNC-6, but lacks UNC-6 functions at longer range. RNA-seq shows that *unc-6(tm)* can encode a transmembrane version of UNC-6 (UNC-6(TM)). Possibly, the VD growth cone is initially polarized with a close-range, potentially contact-mediated interaction with UNC-6, but maintenance of polarity requires a diffusible, long-range role of UNC-6, and membrane anchored UNC-6(TM) cannot provide this long-range function. It is also possible that overall levels of UNC-6 are reduced in *unc-6(tm)*, resulting in this hypomorphic phenotype. However, these results are consistent with *unc-6(tm)* encoding a transmembrane UNC-6 molecule that cannot diffuse at long range.

Consistent with this idea, *unc-6(tm)* and *unc-6(null)* mutants displayed similar levels of AVM and PVM ventral axon guidance defects. AVM and PVM cell bodies reside laterally away from the ventral nerve cord, and thus require longer-range UNC-6, presuming the source is the VA and VB neurons in the ventral nerve cord. This suggests that *unc-6(tm)* is deficient for longer-range roles of UNC-6. Together, these data indicate that a close-range interaction of the VD and DD axons with the *unc-6-*expressing VA and VB motor axons polarizes the growth cone, and that diffusible UNC-6 is required to maintain this polarity as the growth cones move dorsally away from the ventral nerve cord. Thus, UNC-6 might have distinct close-range and long-range guidance functions. A similar close-range interaction might occur in the vertebrate spinal cord, where commissural axon outgrowth requires haptotactic interactions of the axons with the ventricular zone ependymal stem cell processes along the outgrowth path that express Netrin1, and not the Netrin1 source in the floorplate (Dominici *et al.* 2017; Varadarajan and Butler 2017; Yamauchi *et al.* 2017; Morales 2018). Furthermore, *Drosophila* Netrin has short-range effects independent of DCC (Keleman and Dickson 2001). This is consistent with studies in *C. elegans* showing that UNC-5 but not UNC-40/DCC is required for initial VD growth cone polarity (Norris and Lundquist 2011; Mahadik and Lundquist 2023a), and that UNC-40 drives growth cone protrusion in response to UNC-6 (Norris and Lundquist 2011; Norris *et al.* 2014; Gujar *et al.* 2018). In the Polarity/Protrusion model, initial and ongoing VD growth cone polarity relies on the UNC-5 receptor, which also inhibits growth cone protrusion laterally and ventrally. UNC-40/DCC stimulates protrusion dorsally in response to UNC-6, which results in net dorsal growth cone migration. Growth cone size in *unc-6* mutants is not increased as it is in *unc-5* mutants, possible reflecting this dual role of UNC-6 in both stimulating and inhibiting protrusion.

### UNC-6 from a ventral source is not required for dorsal-ventral axon guidance

The data presented here, along with previously reported *unc-6* expression in the VA and VB neurons (Wadsworth *et al.* 1996), suggest that a ventral directional UNC-6 signal might polarize the VD growth cone at short range, and that diffusible UNC-6 is required to maintain polarity. However, *unc-6(+)* expression from ectopic, non-ventral sources nearly completely rescued the VD/DD axon guidance defects and VD growth cone polarity defects of *unc-6(tm)* and surprisingly, *unc-6(null)* mutants. *unc-6(+)* expression from the pharynx driven by the *myo-2* promoter, and expression of *unc-6(+)* from dorsal body wall muscle using the *slt-1* promoter both rescued. It is possible that these transgenes drive expression of *unc-6* in ventral tissues such as VA and VB, although this expression has not been reported. Furthermore, ectopic expression of *unc-6(+)* from these constructs did not significantly affect VD/DD axon guidance or VD growth cone polarity in a wild-type background, indicating that axons cannot be redirected by ectopic UNC-6. Together, these data suggest that a ventral directed source of UNC-6 is not required for dorsal VD/DD axon outgrowth.

Ectopic expression of *unc-6(+)* also strongly rescued ventral AVM/PVM and HSN axon guidance of both *unc-6(tm)* and *unc-6(null)* mutants, suggesting that a ventral directed source of UNC-6 is also not required for ventral axon guidance or the SOAL model of HSN ventral axon guidance. However, *unc-6(+)* expression in a wild-type background from *Pmyo-2* and *Pslt-1* did weakly but significantly interfere with AVM and PVM ventral guidance. This suggests that AVM and PVM might respond differently to UNC-6 than VD/DD, and ectopic UNC-6 might interfere with AVM/PVM guidance.

In the HSN neuron, which extends an axon ventrally to the ventral nerve cord, UNC-40 appears to have an inherent ability to cluster at the cell membrane (Kulkarni *et al.* 2013; Yang *et al.* 2014; Limerick *et al.* 2017). The orientation of these clusters ventrally and subsequent ventral protrusion and extension requires UNC-6, although it is unclear if the UNC-6 source needs to come from a ventral direction. Rescue of HSN ventral guidance defects by ectopic *unc-6* expression suggest that a ventral signal is not required to orient UNC-40 clusters in the HSN in the SOAL model.

Previous studies showed that ectopic *unc-6* expression could redirect or perturb axon guidance, including dorsal-ventral placement of the sublateral nerves, the ventral nerve cord, and SDQL migration (Kim *et al.* 1999; Ren *et al.* 1999). Here, ectopic *unc-*6 expression caused no defects in VD/DD axon guidance and minor defects in AVM/PVM axon guidance. Possibly, different neurons have distinct requirements for the source of UNC-6, or levels and/or timing of expression differed in these experiments. In any event, ectopic *unc-6* expression rescued dorsal-ventral axon guidance defects, suggesting that a ventral source of *unc-6* is not required.

How might UNC-6 mediate directed growth cone outgrowth from a non-directional source? One possibility is that UNC-6 acts as a permissive, rather than instructive, signal for dorsal or ventral guidance, and that responsive cells have inherent dorsal-ventral polarity that is activated by UNC-6. Dorsal-ventral embryonic polarity is established upon the division of the AB blastomere, with the ABp daughter randomly displacing the P2 blastomere to what will become the ventral region of the embryo (Sulston *et al.* 1983). Possibly, all subsequent cell divisions, including those that generate neurons, maintain this dorsal ventral polarity, and nascent neurons are pre-programmed to respond to permissive signals about directional axon outgrowth.

Another possibility is a second signal. While UNC-6 is required for dorsal-ventral outgrowth, another signal might be required in a directional manner to act with UNC-6. In ectopic expression of UNC-6, the guidance role of UNC-6 would only be active in the presence of this second signal. If the second signal was expressed in the same cells as UNC-6 in the ventral nerve cord (*e.g.* VA and VB), then ectopic UNC-6 would only be active where the second signal is expressed. This second signal could be a transmembrane or secreted signaling or basement membrane protein, or it could post-translationally modify UNC-6, such as a glycosylation enzyme or a protease. Indeed, *unc-6* interacts genetically with the basement membrane protein NID-1/Nidogen (Kim and Wadsworth 2000), and the extracellular matrix can modulate *unc-6* signaling, including orientation of UNC-40 clusters in the HSN (Yang *et al.* 2014). Thus, UNC-6 might be asymmetrically localized or activated in discrete regions of the extracellular matrix, obviating the need for a directed UNC-6 source.

In cultured neurons and neuronal explants *in vitro*, Netrin clearly has an instructive role in attraction or repulsion of axons. While this chemotactic mechanism might work in some contexts *in vivo*, work presented here suggests that UNC-6/Netrin could have a permissive role in dorsal ventral VD/DD, AVM/PVM, and HSN axon guidance *in vivo* in *C. elegans.* Ectopic expression of UNC-6/Netrin did not severely perturb dorsal-ventral axon guidance, which would be expected in a chemotactic gradient model. The gradient model implies that growth cones dynamically sense Netrin levels and change their behavior accordingly. For example, growth cones attracted to Netrin would protrude more robustly in the direction of Netrin, and repelled growth cones the opposite. Our results indicate that ectopic UNC-6/Netrin sources cannot redirect axon outgrowth *in vivo*. In the second signal hypothesis, activated UNC-6/Netrin from the ventral nerve cord could form a gradient. However, in *unc-5* and other mutants, extent of growth cone protrusion can be uncoupled from growth cone polarity (Gujar *et al.* 2018; Gujar *et al.* 2019). Large, overly protrusive growth cones can be unpolarized, and small growth cones with reduced protrusion can maintain their polarity. Results presented here are consistent with the Polarity/Protrusion model of directed growth cone migration, wherein UNC-6/Netrin first polarizes the VD growth cone in a short-range or contact-mediated event, and then maintains that polarity at a longer range as the growth cone migrates away from the ventral nerve cord. In both dorsal guidance (the Polarity/Protrusion model) and ventral guidance (the SOAL model), the role of UNC-6 appears to be to polarize the pro-protrusive activity of the UNC-40/DCC receptor in the direction of axon outgrowth.

## Acknowledgements

Thanks to the Lundquist and Ackley labs for discussion and suggestions, to B. Sanderson, of the National Institutes of Health Kansas Infrastructure Network of Biomedical Research Excellence Informatics Core (P20GM103418) and the KU Center for Genomics, for assistance with RNA-seq analysis, and Wormbase. Some *C. elegans* strains were provided by the *Caenorhabditis* Genetics Center, funded by National Institutes of Health (P40 OD010440). High-throughput sequencing was conducted by the KU Genome Sequencing Core, part of the National Institutes of Health *Center for Molecular Analysis of Disease Pathways* (P30GM145499).

